# Dynamics and Internal Control of Body Temperature in Response to Infectious Agents and Other Causal Factors

**DOI:** 10.1101/566679

**Authors:** Charles D. Schaper

## Abstract

Thermoregulation is crucial to homeostasis, but the mechanisms of its dysfunction are still largely mysterious, including fever, which is generally the most disconcerting sign of a serious infection or disease. Theories on body temperature dynamics that aim to explain a fever, such as changes in an internal setpoint, have been proposed, but none can identify the fundamental molecular pathways that produce a fever. Here, potential molecular pathways resultant in fever are identified, modeled, and compared to experimental temperature response data. Based on recent developments made by this lab, which has shown that the pyrogen prostaglandin E2 (PGE2) possesses similar binding affinity as the hormone cortisol (CORT) at the critical ligand binding domain (LBD) of glucocorticoid receptors (GR); molecular modeling, mathematical modeling and a case study for validation is used to indicate that competitive inhibition of CORT by PGE2 as a fundamental reason for dysfunctional dynamics of body temperature, including fever. Comprised of a superposition of proportional and derivative terms of signals representing temperature receptors, CORT concentration, and PGE2 concentration, the internal temperature control model characterizes dynamics associated with the cardiovascular, immune, and neural systems in response to infectious agents, triggering events, and other causal factors. The model is validated by examination of the transient and spectral characteristics of a three-day case history involving temperature trajectories after physical activity protocols in response to a standard vaccination of pneumococcal and influenza species.

## 1 Introduction

A recent case history involved a peculiar temperature response comprised of a half hour of shivering, then a rapid rise in body temperature to fever levels that stabilized for several hours, followed by sweating and a return to baseline temperature; all of which took place starting eight hours after a vaccination for pneumonia and flu, and fifteen minutes after the completion of a ninety-minute routine of physical activity. Furthermore, a similar response, attenuated, was repeated for the subsequent two days before clearing. Of particular interest in this response were the interrelationships of vaccine, physical activity, and fever; moreover, the relation of “chills” preceding fever was also of note.

In researching the cause of this unusual temperature response, the literature indicated that body temperature control variations, that is fevers, due to infectious agents was dictated by a change made to the “temperature setpoint” located in the hypothalamus, by which body temperature was altered to reduce the regulation error, [1, 2]. Alternative theories devised stabilization zones and feedforward loops [3] to characterize temperature response. However, these theories were not satisfactory in examining the case history presented above since these theories could not indicate how physical activity could trigger the temperature response, and thus would seem to deduce it as merely coincidental. Because these theories on body temperature control did not explain the temperature response observed in the case history, a fundamental analysis of body temperature control was pursued.

As the initial step in analyzing the temperature response, a mathematical model based on systems theory was derived that could indicate the model order of the temperature trajectory, permitting the identification of a linear input-output relation, mapping physical activity to temperature trajectory. The model did not propose chemistry, or a molecular foundation, as to the cause of the temperature response, but indicated that the response seemed to involve an interplay between the cardiovascular system and the nervous system, of which the immune system had influence. The cardiovascular system was incorporated in order to explain the aspect of physical activity, and the model had indications that some type of molecule associated with the cardiovascular system was interacting with a molecule of the immune system, which caused deviations in temperature control and an oscillatory response. A model order describing the response was predicted, and the temperature response was simulated well and described in a preprint [4].

To enable a basic understanding of the causative source of fever, the identification of the molecules that were responsible for fever was pursued, of which the starting point of the investigation was the characteristics determined by the mathematical model, namely: an immune system associated molecule that was capable of causing fever, and a cardiovascular system molecule that was a function of physical activity. After a brief evaluation, it became evident that the immune system molecule prostaglandin E2 (PGE2), which is a pyrogen that can be derived from infectious agents, and the cardiovascular system molecule, cortisol (CORT), a hormone, merited evaluation. Subsequent analysis indicated a striking resemblance of the two in chemical elements and molecular weight. Furthermore, a conformer of PGE2 resembled the functional group positioning of cortisol, and could be coordinated with the ligand binding domain (LBD) of glucocorticoid receptors (GR). Molecular modeling indicated that the GR-PGE2 structure was actually at a lower energy state than that of GR-PGE2. The results were documented in the preprint [5].

Here, the newly found association of CORT and PGE2 at the LBD is used to revisit the mathematical model of fever previously derived absent chemistry, although the model predicted its characteristics. Here, material and energy relations are added for CORT and PGE2, including the hypothalamic-pituitary-adrenal (HPA) axis, and the pathways are specified and integrated into the model to analyze temperature and fever. The model is verified by comparing it to the case history of the temperature response including hypothermia and hyperthermia. Thus, a basic framework is developed for thermoregulation, which is vital to homeostasis and to understanding of basic physiology.

Moreover, this study on fevers can be adapted to other signs and symptoms of disease, particularly infections. As the hypothalamus is the central controller of the autonomic nervous system, disruption due to the processing of cortisol via prostaglandins, will cause misprocessing of other functions besides temperature. In a similar manner, other aspects such as fatigue and even emotional dysfunction, such as depression, may be impacted by disease through the molecular pathways described in this article. Some of these issues are described in the discussion section.

## 2 Results

It is the thesis of this article that through the competitive inhibition of cortisol by PGE2 at the ligand binding domain of glucocorticoid receptors, fever is derived. In sum, infectious agents that ultimately result in the production of prostaglandins from the arachidonic acid pathway, yield a situation in which cortisol is not processed according to its designated rate by hypothalamic cells. In turn, this results in altered transient behavior of both the resultant neural and neuroendocrine signals of the autonomic temperature control system, producing deviations in body temperature from normative values. This section describes the site and pathway of competitive inhibition of CORT by PGE2, and develops a linearized set of equations as a mathematical model describing the deviations in body temperature from infectious agents and other causal factors. The mathematical model is then evaluated in a case study of fever and associated chills as generated by a vaccination, triggered by the abrupt cessation of physical activity.

### 2.1 Molecular Pathways Resultant in Fever

The basic molecular pathway comprising the underlying mechanism resultant in fever is developed in this section, and is presented in Figure 1. The source of biomaterials to this pathway include arachidonic acid (AA), which can be obtained from lipopolysaccharides (LPS), such as with gram-negative bacteria, and from the phospholipid bilayer. The enzyme family, cyclooxygenease (COX), can be obtained starting from the cytokines, including the interleukin materials, and TNF-α. From the cytokines, the NF-κB and NFAT pathways produce COX [6, 7, 8, 9]. The combination of AA and COX will generate PGE2, at a rate that will be limited either by the source material of AA or the enzyme COX. For example, gram negative bacteria produces LPS and thus result in an increase in AA; on the other hand, gram positive bacteria, which does not contain LPS, can enter the pathway by increases in TNF-α, which will increase COX, and improve the conversion rate of any AA available, or provide a driving force for the creation of additional AA.

**Figure 1:**
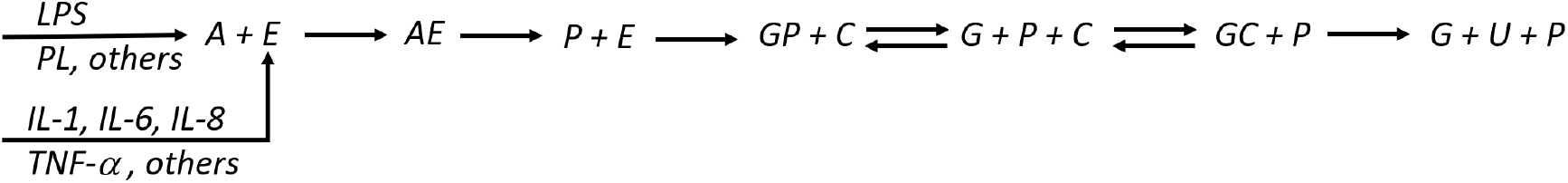
The pathways of producing body temperature variations (and other signs and symptoms of disease). The source streams to the system are associated with the formation of arachidonic acid (A) and the enzyme cyclooxygenase (E), which can generated through lipopolysaccharides, from gram negative bacteria, or simply the phospholipid cell membrane. The enzyme can be derived through for example the cytokines, which can result in cyclooxygenase through the nf-κB pathway. The combination of A and E can complex and result in prostaglandin E2 (P). Then, P makes its way to the LBD of the glucocorticoid receptor (G) which nominally interacts with cortisol (C). Through competitive inhibition, eventually C is established with G for sufficient time to permit the formation of the intended cellular response (U).

After the formation of PGE2, it will compete with CORT for the LBD of the GR, which this lab initially identified [5]. When PGE2 occupies the GR, the PGE2-GR complex will not function as does CORT-GR, but since the PGE2-GR is relatively stable compared with CORT-GR, the percentage of time that it occupies the LBD will influence the overall kinetics of the pathway in converting the normative response of CORT-GR. When PGE2 is replaced in the LBD by CORT, the complex becomes active, and after expression results in the normative response from CORT, in the figure represented by the variable *U*.

Through molecular models, it has been shown in [5] that PGE2 can be conformed such that it and cortisol (CORT) will have a similar arrangement of chemical elements and functional groups which would permit equivalent affinity with the LBD of GR. Cells containing GR are found in almost every cell and tissue, but should be particularly relevant to the hypothalamus that is the control site for temperature, and the site for control of the HPA axis in response to cortisol. The association of PGE2 and CORT is compared in Figure 2, which represents the electrostatic potential, molecular dimension and positioning of the structures of Cortisol and a conformational isomer of PGE2 at the LBD. The five amino acid residues comprising the LBD are defined in this figure, and the position of alpha carbons are actually very similar, within 2 Å.

**Figure 2:**
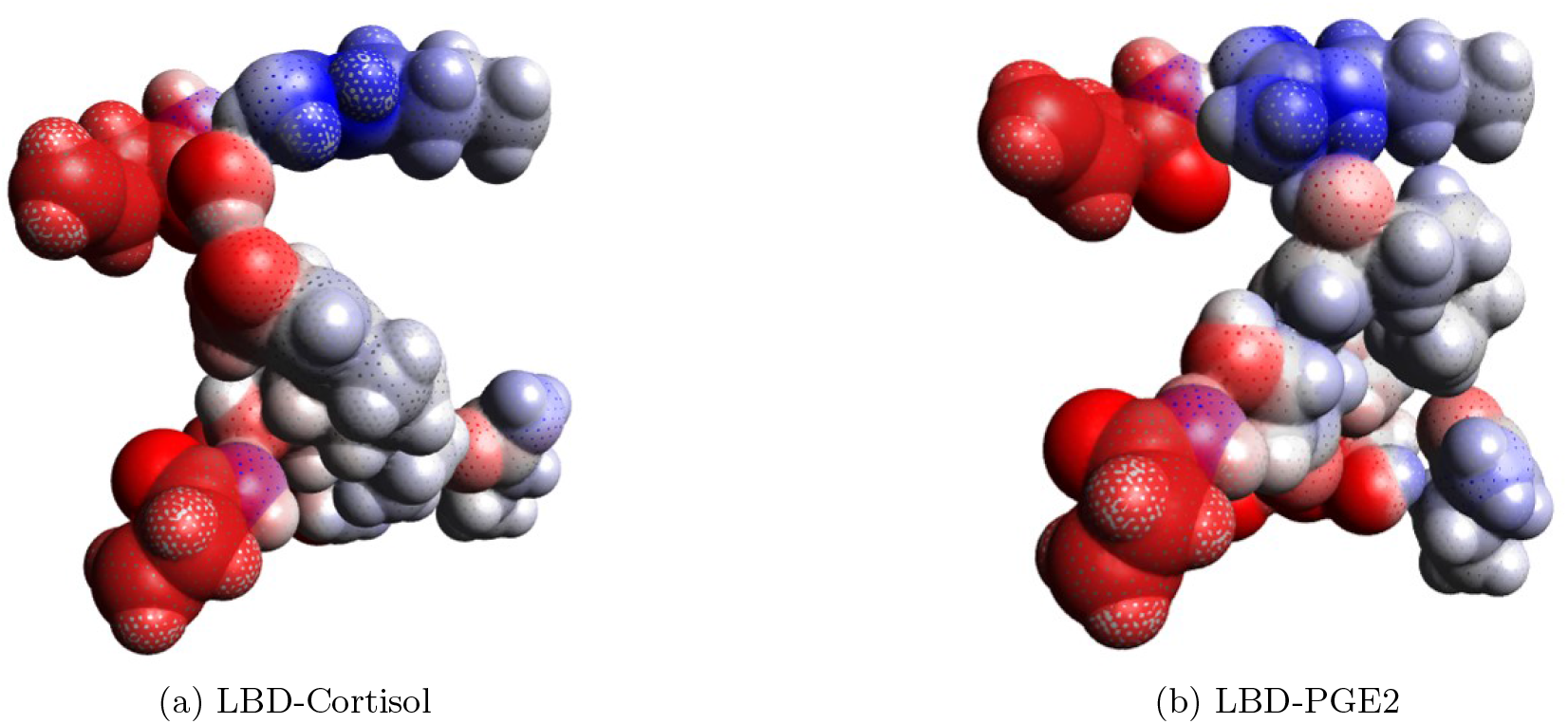
A comparison of the interaction of (a) CORT and (b) PGE2 of electrostatic potential when configured with the ligand binding domain of the glucocorticoid receptor. Visually, the results appear similar in terms of the orientation of the alpha-carbon positions of the residues of the amino acids comprising the LBD, and the overall distribution of electric charge. In determining the spacing between the residues, all are less than 2 Å difference in positioning.

The energy state associated with PGE2 at the LBD, both with and without calcium ions, is lower than CORT, thus indicating that PGE2 is relatively more stable, although it is smaller relative to CORT by 33% when calcium ions are integrated as shown in Figure 3. Thus this indicates that the conformational shift of the LBD is different between PGE2 and CORT. Moreover, depending upon the kinetics, the rate of this process of calcium ion introduction is influential of the cellular response. That is, if the concentration of CORT within the cytosol impacts the introduction of new CORT molecules into the cell through the phospholipid bilayer it will produce a forcing function as a concentration gradient. It indicates that the action of the neurotransmitter to the cell will be influenced by the concentration gradient induced by the local variations in the concentration of cortisol and PGE2 within the cell, since they are associated with a calcium flux and depolarization. These concepts are developed further in the modeling section.

**Figure 3:**
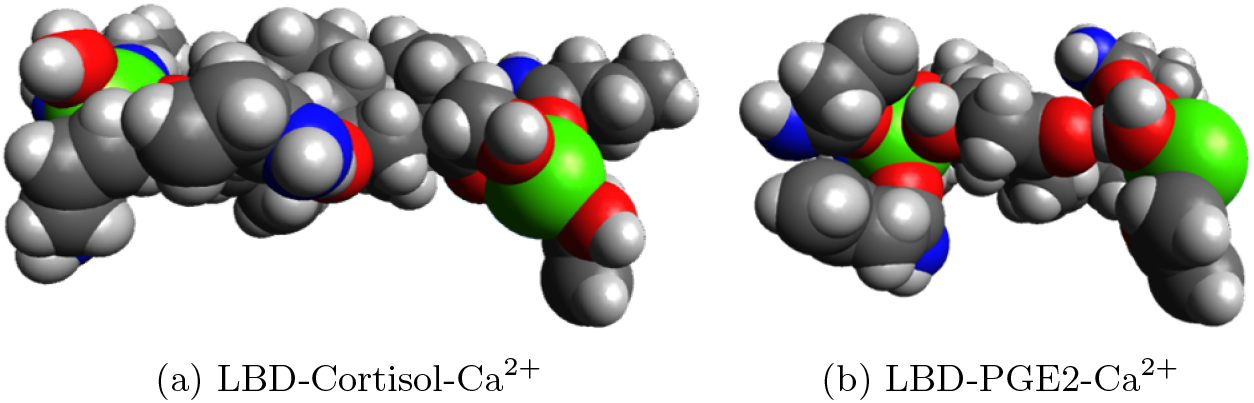
Without the addition of calcium ions, the configuration of CORT and PGE2 with the LBD is quite similar in positional configuration. However, with the addition of calcium at the areas of high electrostatic potential, there is significant difference in the positions of the residues comprising the LBD of (a) CORT and (b) PGE2. Thus, the expression profiles will be different, although the energy associated with PGE2 is less than that of CORT, and therefore increased stability.

### 2.2 System Modeling of Body Temperature

To develop a model that can capture the dynamics associated with the production of a fever, pursuant to the influence of CORT and the competitive inhibition by PGE2 at GR, a mathematical model is developed of body temperature dynamics. While the modelling approach is general enough to incorporate other signs and symptoms of disease pertaining to the competitive interference at the GR by CORT and PGE2, the focus in this section, as it is in the article, is on body temperature. The resultant mathematical model is a lumped parameter structure whose parameters can be identified from input-output data and validated by its analysis and predictive capability, as well as its internal structure. An important aspect of this analysis is the primary feedback control law developed for the hypothalamus, as it will determine its overall integration and coordination within the network of systems, including the cardiovascular and the nervous systems, and its interaction with causal factors, including infectious agents and internal stimulation. From a high level, the model is a neural and neuroendocrine mapping of infectious agents to body temperature dynamics and control.

## 3 Temperature Control Policy

In addition to the newly developed molecular pathways characterizing fever, another innovation of this research article is the temperature control laws presented in this section which utilize not only feedback from temperature, but also from the hormone cortisol to define the neural component of temperature control. To quantify such dynamics, a control law mechanism is developed by which the input signal sent to the target cells from the hypothalamus is generated in response to neural inputs from temperature receptors in conjunction with neuroendocrine inputs via cortisol signaling, which act to depolarize the cell through the addition of calcium ions. Moreover, the temperature control policy is expressed as a function not only of the concentration of cortisol, but also as the rate of change. The rationale for the incorporation of the time derivative of cortisol is in the control law expression is because the transport of cortisol into the cell also associates the ionic transport of calcium, leading to an accumulation of charge about the phospholipid bilayer, and depolarization of the cell. This essentially establishes a localized concentration gradient of cortisol within the cell itself, and the potential for build-up of material within the cytosol, leading to a variable diffusion gradient for the flow of cortisol into the cell, and thus an altered rate of change of cortisol concentration. Hence, in an integration with the neural signals associated with temperature and its rate of change and the concentration of cortisol and its rate of change, the temperature control law, by which temperature compensation signals are made, follows by modeling the electric charge balance expressed as a functional relation:

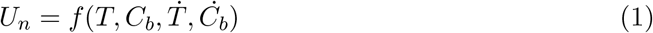

where *U_n_* is the neural action potential as an input signal to the internal temperature control system that emanates about the boundary of the nerve cell membrane, *T* denotes an effective temperature associated with the body, *C_b_* is the concentration of cortisol at the boundary of the membrane, 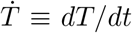 and 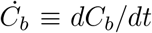, which are the rates of change associated with the temperature and boundary cortisol concentrations.

For the control law expression, the effective temperature *T* is considered as a thermoreceptor signal that is representative of body temperature, which can be obtained from a combination of thermoreceptors and averaged over time. As noted, the input signal computed, *U_n_*, is modeled here as a just a scalar to the temperature control system. It is considered to be distributed in an appropriate manner to the various vector of mechanisms of heating and cooling, such as metabolic energy generation, vasodilation, as well as sweating and shivering. For the purposes of this study, the distribution of the output signal to the effector mechanisms is secondary to the analysis as it will not impact the present analysis, and can be considered as an averaged input. Following these approximations, to obtain a mathematical representation of the control law, a Taylor Series expansion [10] of Equation (1) is taken about a nominal operating point denoted as *U*_0_, *T*_0_, *C*_0_, which would be associated with a standard body temperature, and truncated after the first term results in the expression,

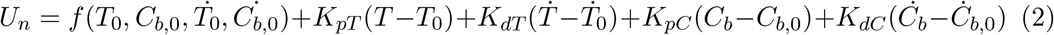

of which the control law scaling factors, which are critical in its performance, are given by:

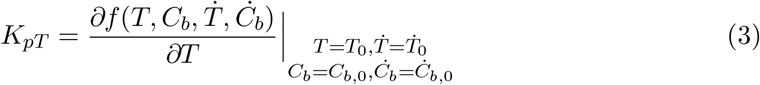

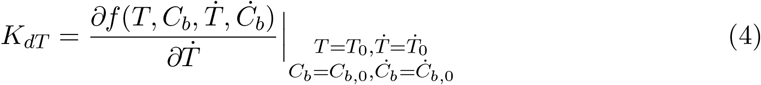

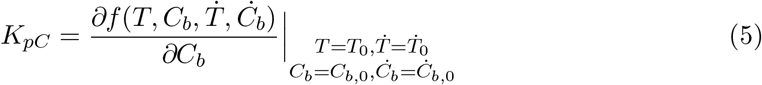

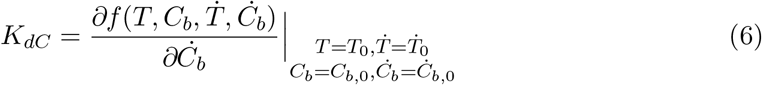

Deviation variables are defined as *ΔU_n_* = *U_n_* – *U*_*n*,0_, *ΔT* = *T* – *T*_0_, *ΔC_b_ = C_b_* – *C*_*b*,0_, and thus we have the control law expressed after substitution into Equation (2), assuming steady-state conditions with 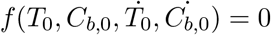, as

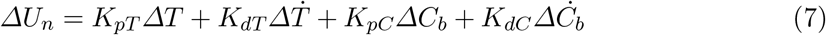

It is noted that the control law, Equation (7), is considered as having a PD (Proportional Derivative) form, which is deployed in industrial process control with success. The proportional terms provide a means of using a feedback control mechanism to manipulate the input variable so as to steer the controlled signals about a nominal value. The derivative terms is primarily to shape the response to desirable trajectories, for example speeding-up the control action. Further, it is noted that the assumption of steady-state conditions is developed in this model, which is standard analysis technique in process control. However, if it is not steady-state, then the term would be included in the control law, and carried through the calculations to have the impact of an impulse disturbance to the system that would be resolved through closed-loop control.

As indicated, one of the key contributions of this analysis is the incorporation of a feedback mechanism based on cortisol. Besides priming the temperature response, it also incorporates a signal in the control law that is representative of the status of the response from the input signal, which could be, for example, that result in internal metabolic generation as regulation of temperature, which would alter the cortisol concentrations. Moreover, based on observable responses to infectious agents, it appears mandatory to incorporate such a coupling mechanisms, since a feedback controller that just fed back on temperature would not be able to capture the impact of infectious agents as detailed in the validation example of Section 3.8.

### 3.1 Corticosteroid releasing hormone

In producing the control signal associated with internal temperature variations, the cortisol concentrations to the hypothalamus will also influence the release of additional cortisol through the HPA (hypothalmic-pituatary-adrenal) axis, initially by the formation of CRH (corticosteroid releasing hormone). This is included in the model because the regulation of cortisol will impact body temperature, both as a functional input to the hypothalamic area in controlling temperature, and at the target tissues. The control function expressing the concentration of CRH, denoted as *C_CRH_*, is modeled as:

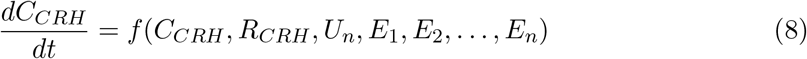

where *R(CRH)* is the reaction rate associated with producing an expressive response, and the exogeneous inputs, [*E*_1_, *E*_2_, …, *E_n_*] that can have effect on body temperature, such as stressors, including those of a physical or cerebral nature. After Taylor Series expansion about an initial condition, we thus have

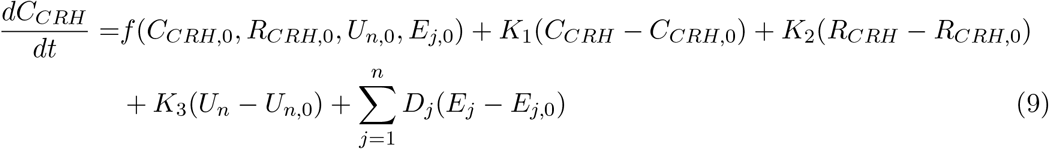

of which the control law scaling factors are given by:

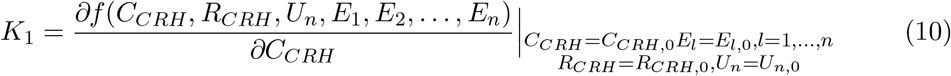

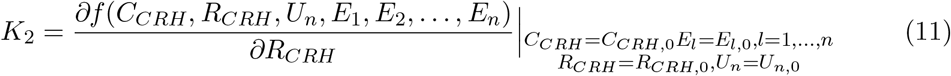

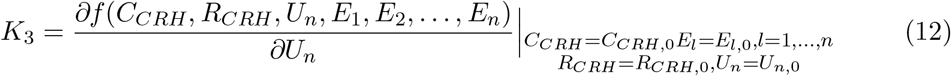

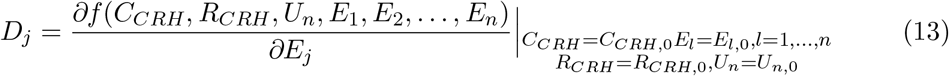

Deviation variables are implemented and with *f* (*C*_*CRH*,0_, *R*_*CRH*,0_, *U*_*n*,0_, *E*_*j*,0_) = 0 at steady-state conditions, the expression becomes:

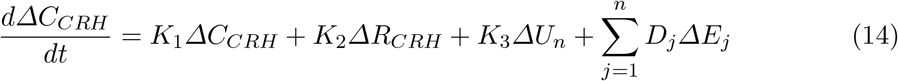

### 3.2 ACTH concentration

Through the HPA axis, CRH will initiate the response of adrenocorticotropic hormone (ACTH), which can be expressed as:

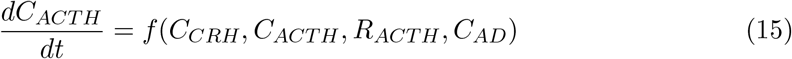

where *R_ACTH_* is the reaction rate in producing an ACTH and in competitive inhibition with *P_ACTH_*, the concentration of PGE2, and *C_ACTH_* is the concentration of ACTH, and *C_AD_* is the concentration of cortisol from the adrenal gland. And thus, using the similar Taylor Series expression, we have:

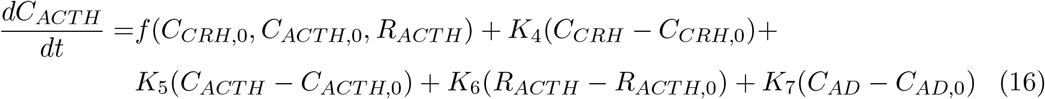

of which the control law scaling factors are given by:

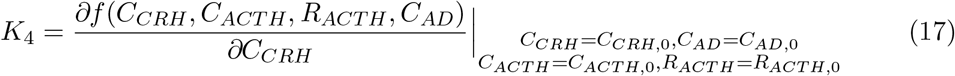

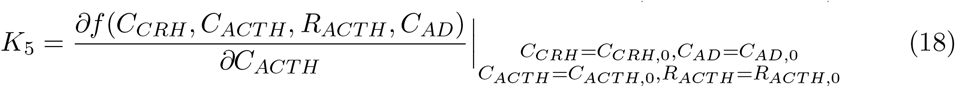

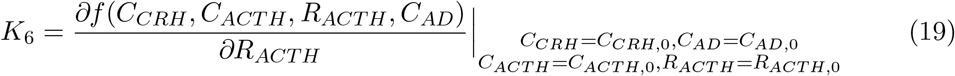

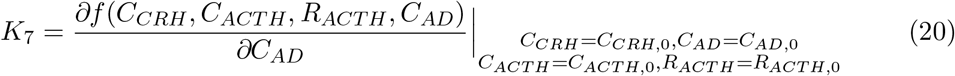

After expression in deviation variable, we have, similar to that for CRH:

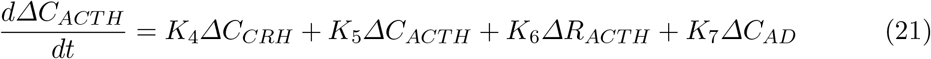

#### 3.2.1 Cortisol concentration from the adrenal gland

The adrenal gland output from stimulation by ACTH can be considered in a similar fashion as that of ACTH, namely:

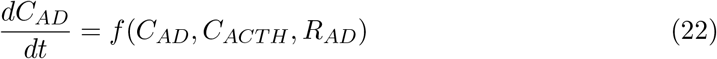

where *R_AD_* is the reaction rate at the adrenal gland involving the glucocorticoid recepor. Following a linear approximation, we have:

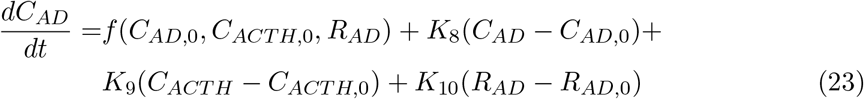

of which the control law scaling factors are given by:

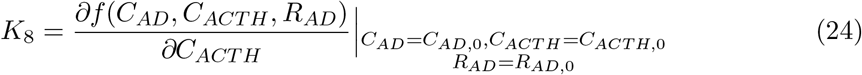

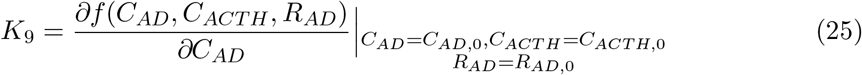

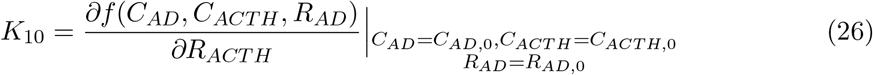

Deviation variables are implemented, and thus,

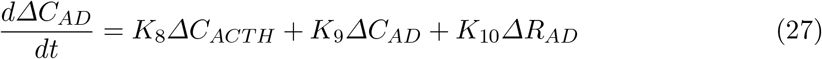

with initial steady state conditions taken as zero.

### 3.3 Cortisol Circulating

In this article, in the development of the feedback control law of temperature, it is proposed that cortisol, which is indicative of the status of the effects of the controller output signal in its response to temperature. It is further proposed that cortisol interferes with external agents, such as infectious components or vaccines, and whose rate of association is designated as *R_b_*, and thus the infectious agent interferes with the temperature control signal. In the development of the model, also incorporated is the potential for an external feed stream *F* that can provide an external source of cortisol. The resulting function is expressed therefore as follows:

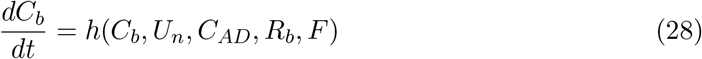

and in producing a linearized representation of Equation (28), a Taylor Series expansion truncated after the first term yields:

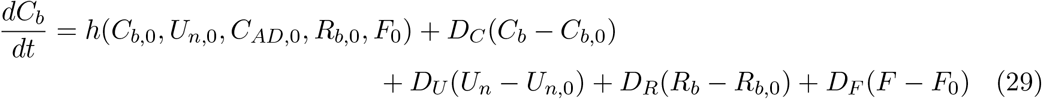

where

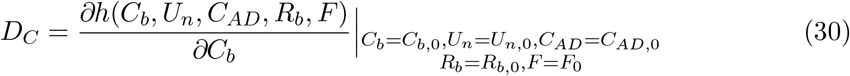

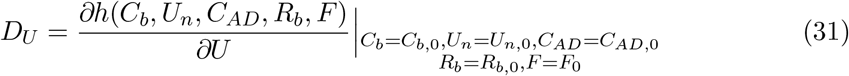

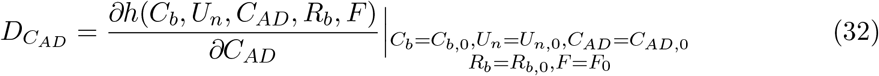

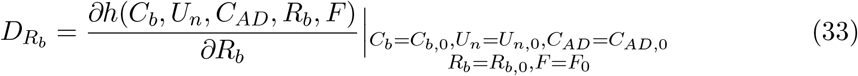

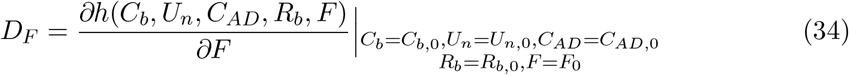

And substitution and then utilizing deviation variables *ΔF* = *F* — *F*_0_ and *ΔR_b_* = *R_b_* — *R*_*b*,0_, the following equation for cortisol circulating is obtained,

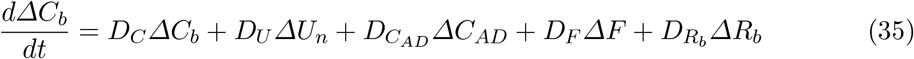

### 3.4 Hypothalamic Cortisol

For the intracellular cortisol concentration, denoted as *C*, at the decision point in the hypothalamus of producing the feedback control signals based on temperature receptor signals, we have

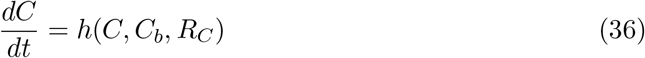

and in producing a linearized representation of Equation (36), a Taylor Series expansion truncated after the first term yields:

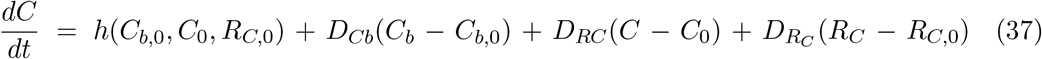

where

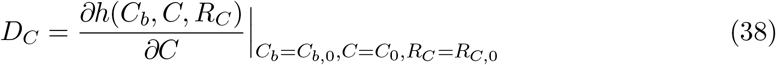

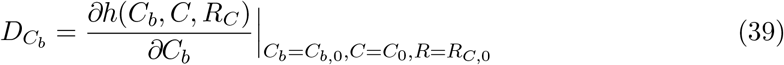

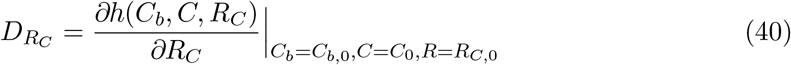

And in using deviation variables with zero steady state conditions, we have

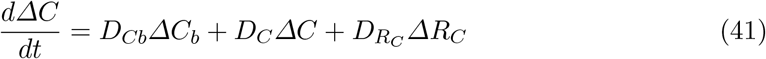

### 3.5 Body Temperature Dynamics

The temperature by which control signals are generated according to the control law of Equation (27) corresponds to an effective signal that is expressed as a function shown in Equation (42), which is comprised of not only variables *T* and *U*, but also exogeneous inputs, [*E*_1_, *E*_2_, …, *E_n_*] that can have effect on body temperature, such as exercise, environmental factors of humidity and temperature, external heating, and other inputs including stressors such as those associated with deviations of the sympathetic and parasympathetic nervous system, and thus *E_j_* (*s*) can incorporate inputs due to emotional stress, cerebral thoughts that may induce sympathetic activity, torpor, hibernation and other activities. Open-loop expressions of temperature models, which are first order, and its influence from external factors such as drugs and exercise have been developed [11]. In the approach developed in this paper, the representation for temperature is developed in closed-loop format for general sets of inputs. As the body temperature is in general non-uniform, the temperature of concern here is an “effective” temperature, that is one in which the control laws ultimately respond after integration of the neurological thermoreceptors. The effective temperature dynamics are expressed as

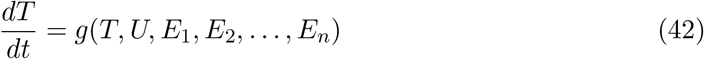

A linear dynamic model is obtained from a Taylor Series expansion of Equation (42) and then neglecting all terms of order two and higher, to arrive at the expression:

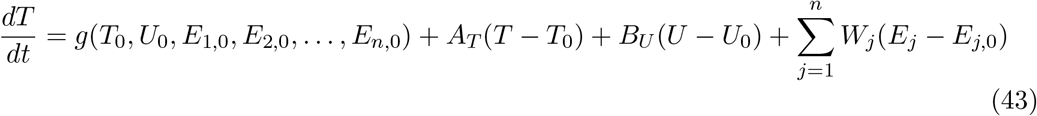

where the model parameters are found from:

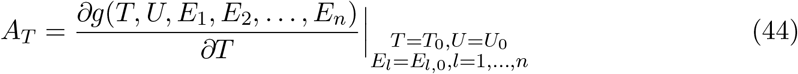

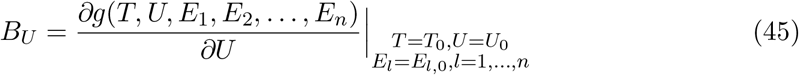

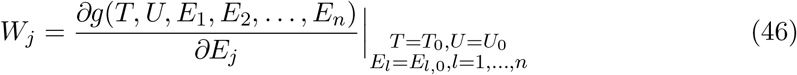

Utilizing deviation variable format with *ΔE_j_ = E_j_ — E*_0,*j*_ about a nominal level, which could be 0, an expression for the temperature dynamics is achieved:

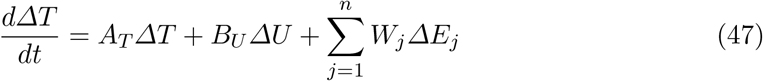

Equation (47) is a linearized model which maps the temperature state trajectories, inputs from the control law, and exogeneous inputs to changes in temperature, and is typical of the dynamic response of thermal systems when used for control system analysis about nominal operating windows. And where the total input to the temperature control system is determined as

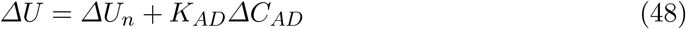

to reflect the neurological and the neuroendocrine input to the system.

### 3.6 PGE2 dynamic model

For the circulatingPGE2 *P_b_*, its dynamic model is represented from a species balance as follows:

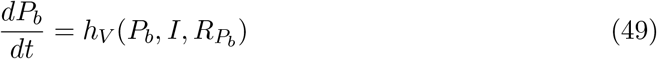

where *I* is the mechanism by which the prostaglandin ultimately enters the system, such as with an injection. The model is also a function of the association rate *R_b_* with cortisol. A Taylor Series expansion truncated after the first term yields,

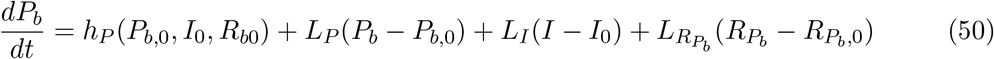

where

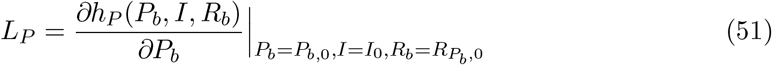

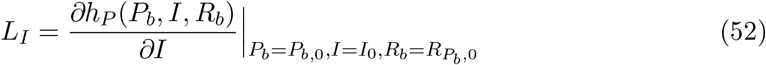

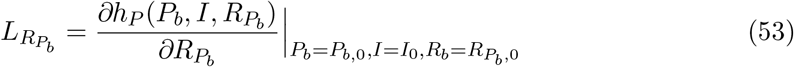

Taking deviation variables, *ΔI* = *I* — *I*_0_, yields

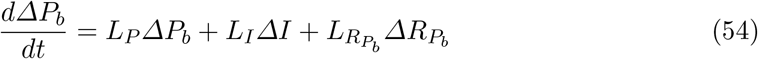

Similarly, for the concentration of PGE2 within the hypothalamus, we have

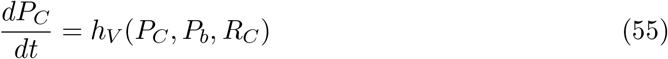

A Taylor Series expansion truncated after the first term yields,

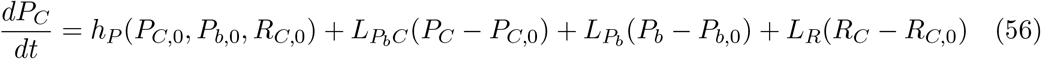

where

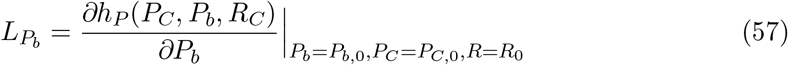

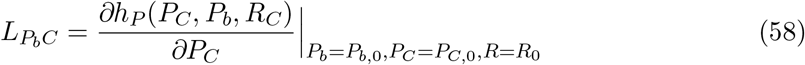

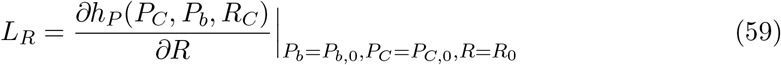

Taking deviation variables yields at zero steady-state conditions

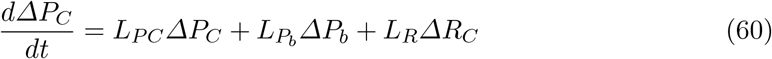

To complete the model for the prostaglandins, using a similar format for the HPA axis, we have the following for the concentration at the other zones of the HPA axis:

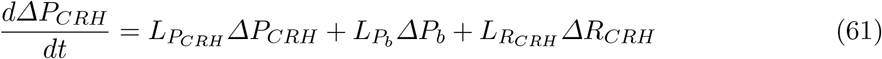

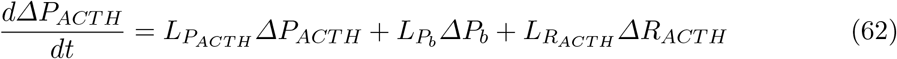

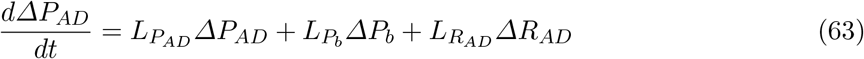

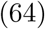

### 3.7 Competitive Inhibition

For competitive inhibition of cortisol and PGE2 within the hypothalamus, the rate can be determined through the following functional:

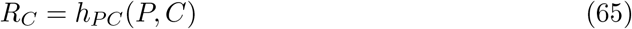

Linearizing the reaction rate representation of Equation (65) yields:

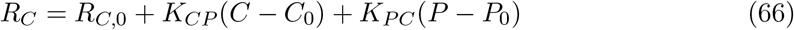

where

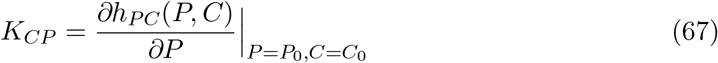

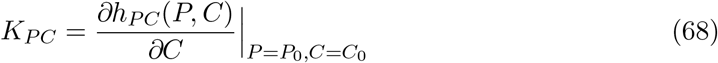

and thus,

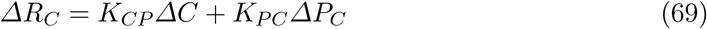

Similarly, for the association rate within the circulation outside of the central nervous system *R_b_*, we consider kinetics which are dependent upon the concentration of PGE2 and the concentration of cortisol:

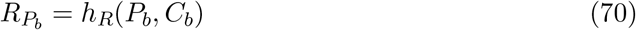

While the rate coefficients may be a function of temperature, it is considered that such effects will be secondary and thus can be neglected without loss of generality. Linearizing the reaction rate representation of Equation (70) yields:

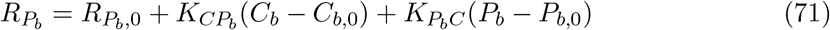

where

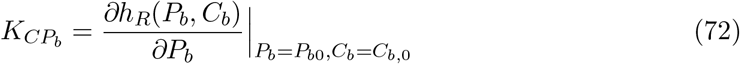

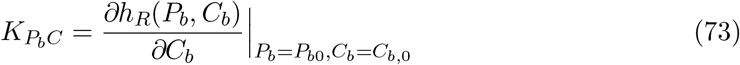

which results in the following expression

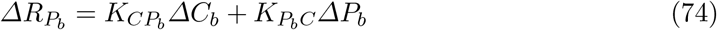

Similarly, the association rates can be determined at the other sites of interaction as follows,

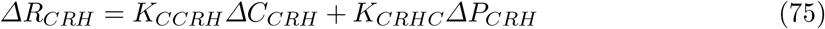

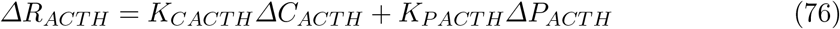

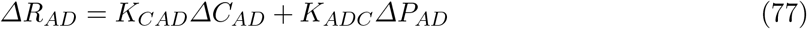

#### 3.7.1 Model Order Reduction

In this section a reduced dimension of the model is presented with several assumptions are deployed: (1) The concentration of the infectious agents causing the disruption in PGE2 concentrations is distributed uniformly throughout the body system, which includes equivalent transient distribution and thus can be represented by a single variable denoted as, *P*, of which the largest association with cortisol is through that associated with *C_b_*; (2) Including the HPA axis and circulatory system outside of the decision center, the hypothalamus, the deviations in cortisol concentration associated with the HPA axis and the resultant concentration within the body are expressed as proportional plus delayed in time, *θ*, and thus the dynamics associated with the circulatory system can be expressed by the single variable as input to the circulating cortisol concentration. The model order for this relation is five in which there are four poles, plus the time delay which can be approximated by a pole-zero Pade term. Additional approximations including that the difference between the cortisol concentration within the circulatory system *C_b_* and at the decision making point for temperature control, *C*, is represented as proportional to the circulatory system and with the time delay which can further be combined with the time delay of the HPA axis, *θ*. In forming a set of linear, time-invariant models to describe the closed-loop temperature dynamics, the LaPlace transform of the modeling equations to develop a lumped parameter model as:

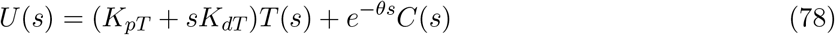

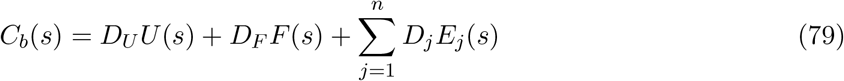

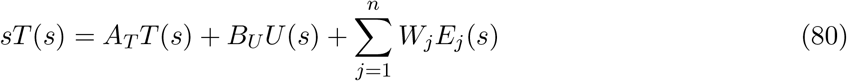

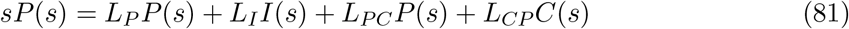

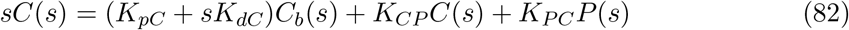

Additional approximations are employed in considering the time delay term associated with the HPA axis generation of cortisol to also incorporate the transportation and mixing delay associated with the distribution within the target tissues, and the concentration of prostaglandin approximately equivalent within the circulation as in the hypothalamus, that is *P_b_*(*s*) ~ *P_C_*(*s*). Due to time periods involving transportation, gene transcription, as well as physical mixing of the materials under evaluation, it is necessary to incorporate time delays into the model. This is important because it will limit the performance of the temperature control system. If there were no time delays, simple high gain controllers could be deployed just around temperature that could minimize the relatively slow variations in temperature due to infectious diseases. However, because the requirements to implement temperature control through the implementation of the endocrine system through metabolic activities, it is necessary to consider such delays. In this model, three time delays are considered: implementation of the CRH pathway; distribution of cortisol throughout the body and into the hypothalamus; implementation of the temperature control command center to the return of the change sensed by the thermoreceptor. A simplified block diagram utilizing the time delay format assuming that the prostaglandin concentration is approximately constant over a period of time, for example the duration of a response. To arrive at the final set of equations to describe the dynamic response and internal control operations of body temperature are taken around a nominal operating point and comprise a set of linear time invarient models. With these approximations and using the pade approximation of the time delay, the closed-loop schematic representation of temperature control is presented in the block diagram of Figure 4, which maps the inputs to the system *E*(*s*), *I*(*s*), *F*(*s*) to the output temperature signal *T*(*s*), m and includes the inner concentration of cortisol *C*(*s*) and PGE2 *P*(*s*). The internal control loop is highlighted indicating the intersection of two coordinated control loops, which will be discussed further in Section 4.

**Figure 4:**
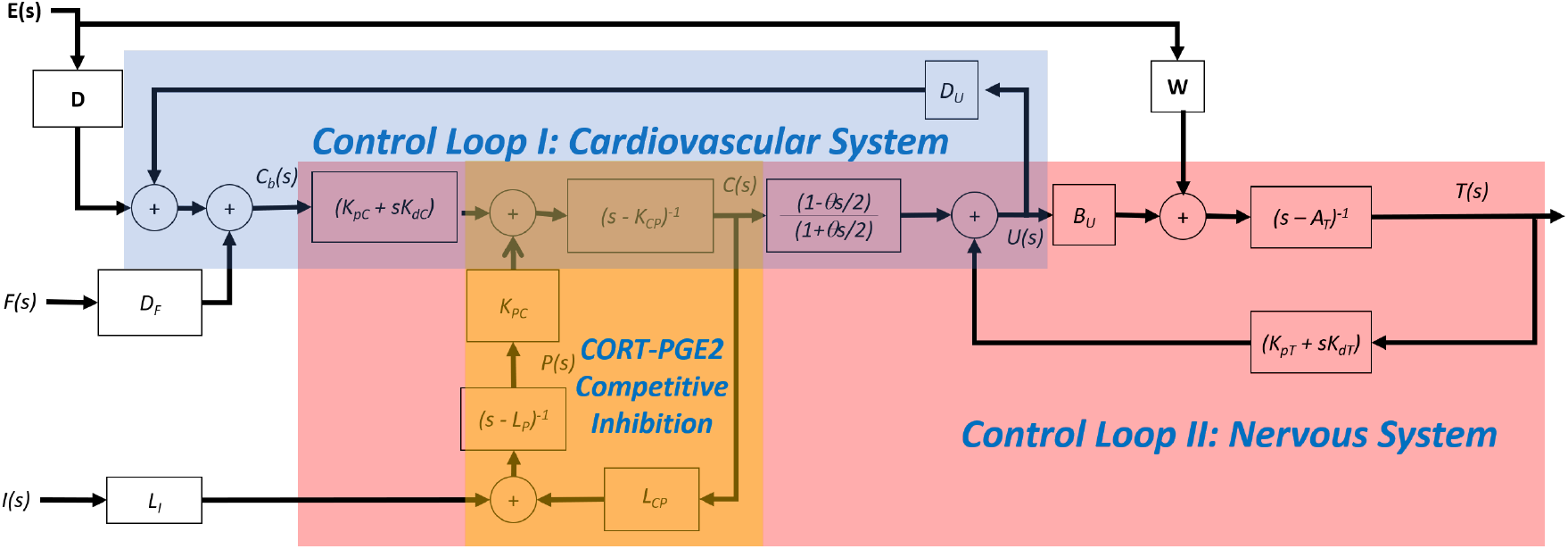
The block diagram of the closed-loop response to temperature of a reduced-order model in response to inputs from infectious agents I(*s*) that results in PGE2 P(*s*) from external pyrogenic agents F(s), and from a vector of external sources, **E**(*s*), which comprise physical activity as well as other external stressors and temperature inducing behavioral factors. The blocks comprise the transfer functions of the model, and include the control law function for U(*s*) which comprises feedback from body temperature T(*s*) and cortisol C(*s*), which is in competitive inhibition with PGE2. This block diagram partitions into sectors demonstrating the integration of the nervous, cardiovascular and immune systems.

It is also possible to obtain a reduced order model by considering instead the competitive inhibition within the target tissues and circulating system outside of the central nervous system. In this case, the input signal for temperature control is immediately obtained at the hypothalamus, and the time delay is taken as due to transportation delay. The inhibition of CORT due to PGE2 takes place in the target tissues, and thus the relations can be expressed as the following set of equations and its block diagram is depicted in Figure 5.

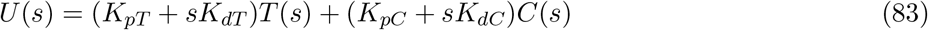

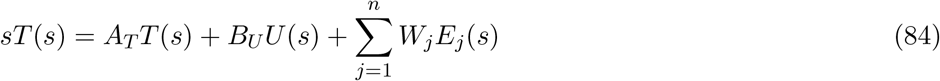

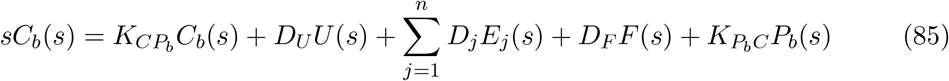

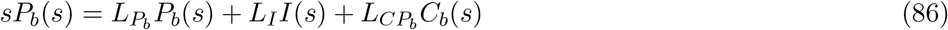

**Figure 5:**
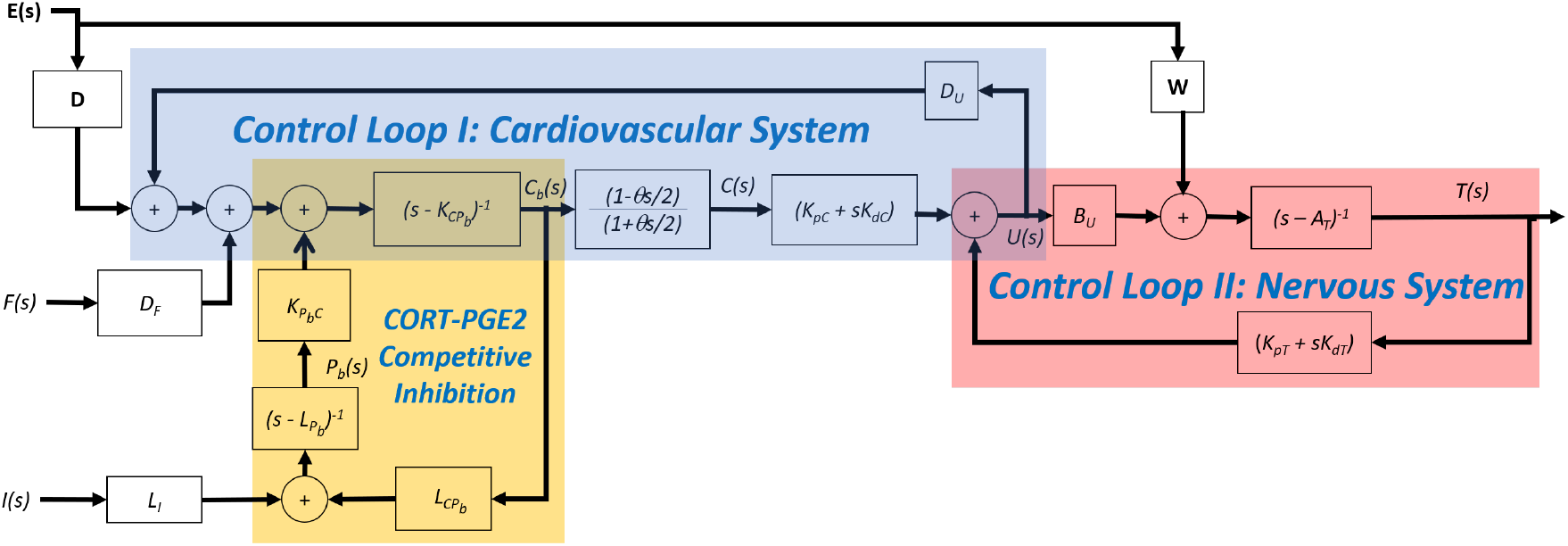
The block diagram of the closed-loop response to temperature of a reduced-order model in response to inputs from infectious agents I(*s*) that results in PGE2 P(*s*) from external pyrogenic agents F(*s*), and from a vector of external sources, **E**(*s*), which comprise physical activity as well as other external stressors and temperature inducing behavioral factors. The blocks comprise the transfer functions of the model, and include the control law function for U(*s*) which comprises feedback from body temperature T(*s*) and cortisol C(*s*), which is in competitive inhibition with PGE2. This block diagram partitions into sectors demonstrating the integration of the nervous, cardiovascular and immune systems.

#### 3.7.2 LTI Models at Multiple Operating Points

In the development of this approach, the output units of interest in characterizing the response are represented as follows:

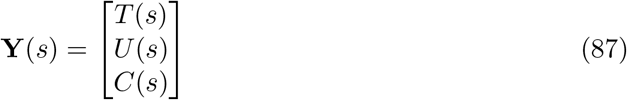

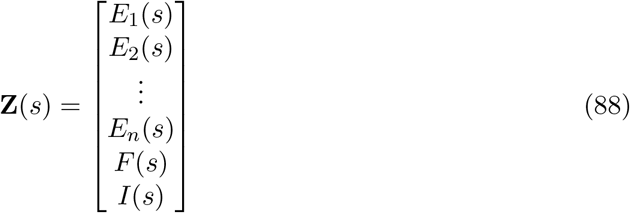

At the linearization point denoted as *k* = 1, 2,…, *r* for a series of *r* LTI models in which an indvidual operating point is given by:

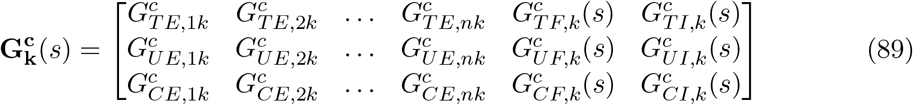

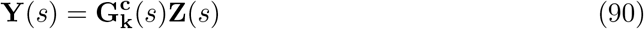

and the closed-loop matrix are specified as,

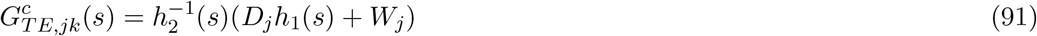

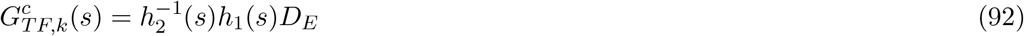

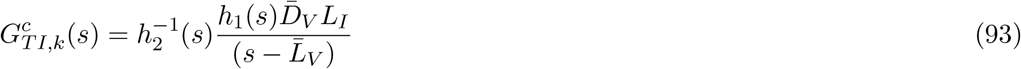

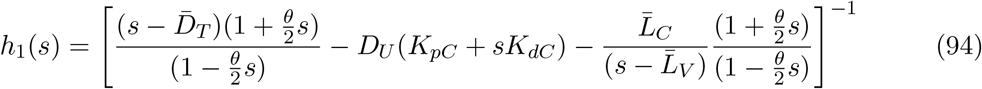

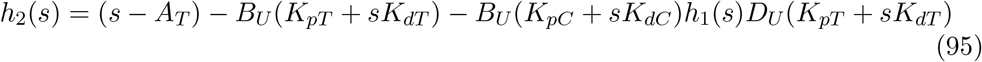

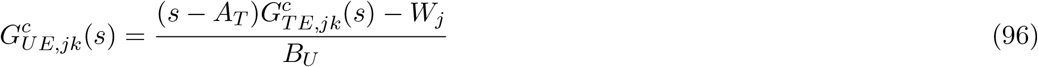

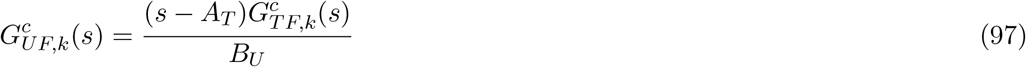

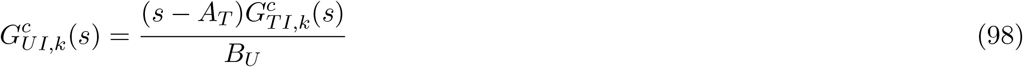

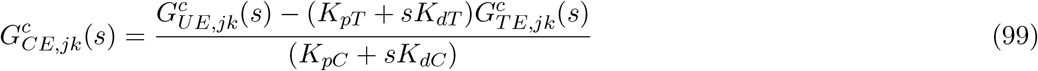

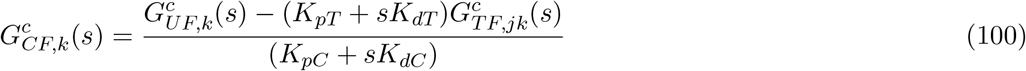

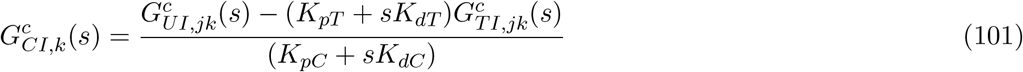

The set of model equations describing the temperature control system about operating points **G^c^**(*s*) is thus represented as

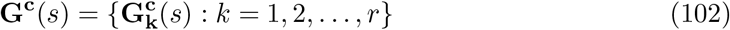

A interesting consequence of the proposed control law mechanism is an intersection of signals associated with the cardiovascular system and with the nervous system linked through the hormone CORT, and includes the immune system response which is in competitive inhibition with CORT at the GR within both the hypothalamus and target organs. The intersection is indicated in Figure 4 with the disturbances induced from external sources, **E**(*s*). The inputs to the control law, whose location is the hypothalamus region, are chemical and electrical in nature, and the output of the control law is neurological and neuroendocrine. Note that the block diagram, which compartmentalizes the immune response with the cardiovascular and nervous system, is formulated as a graphical representation of the mathematical equations describing the process. The control loops are cooperative in the sense that stressors placed on the cardiovascular system, such as increased oxygen demand, can be balanced by the requisite body system response, such as adjustments in body temperature. And conversely, stressors placed on the nervous system, such as the need to dissipate heat, can be balanced by the circulatory response, such as with vasomotor activity. Conversely, the control loops are not run independently, in that changes made to the cardiovascular system are conducted in conjunction with regulation of temperature through the nervous system and thus other body function systems.

### 3.8 Case File

The following case file is analyzed: A participant in reported excellent health, The following case file is analyzed: A participant in reported excellent health (Pulse rate: 45 bpm; Weight 181 pounds; Waist-to-Height Ratio 0.44; Blood Pressure 123/72; Oral Temperature 97.5°F; Cholesterol 128 mg/dL (Standard Range ≤ 239 mg/dL); LDL 29 mg/dL (≤ 159 mg/dL); HDL 74 mg/dL (≥ 40 mg/dL); WBC Count 5.0 K/uL (3.7 – 11.1 K/uL); RBC Count 4.36 M/uL (4.10 – 5.70 M/uL); Hematocrit 41.7% (39.0 – 51.0%); Hgb (14.6 g/dL (13.0 – 17.0 g/dL); MCV 96 fL (80 – 100 fL); RDW, RBC (11.9% (12.0 – 16.5%), Platelets 172 K/uL (140 – 400 K/uL), RBC’s nucleated 0/100 WC) was administered by injection a vaccination associated with pneumococcal and influenza agents at approximately noon. The participant’s temperature was reported as constant prior and immediately after the vaccination shots, and throughout a two-hour period of physical activity (exercise) that took place six hours after the injections. This routine of physical routine had been replicated daily for approximately thirty days preceding the vaccination. Approximately ten minutes after completing the physical activity, which consisted of moderate training with weights and a seven-mile run on a treadmill at an average of 4.6 mile/hour, the participant reported the sensation of severe chills that lasted approximately forty minutes followed by a rapid rise in temperature to a nominal fever, which lasted two hours, followed by sweats in a return to a baseline temperature, thereby concluding the episode seven hours after the completion of the exercise period. The participant reported feeling fine after the episode of temperature fluctuation. On Day 2 and on Day 3, following the same protocol of physical activity at the same time of day, the participant reported a similar response in temperature, through less severe, of which the intensity of the response was estimated as level 8, 3, and 1 for each of the three days, and the period of response was longer during the shivering period but shorter in time at the higher temperatures. The first day required bed rest during the seven hours of high temperature excursion to deal with the sensation of hypothermia and fever, while the second and third day did not require bed rest. By the fourth day, no adverse temperature effects were felt following the protocol of physical activity. To quantify the signs, a numerical assignment is made to the observations: —0.6°F as a sign of significant shivering to 3.2 °F for a fever with a sign of sweating. It was noted that for the pneumococcal vaccine, similar experiences of hypothermia followed by fever were reported from on-line patient feedback, hence the display of temperature effects following vaccination was deemed to be fairly typical, although this case was particularly well characterized as the physical activity regimen was a precursor to a significant dynamic temperature response. The response is characterized in Table 1 to quantify the signs. It was noted that for the pneumococcal vaccine, similar experiences of hypothermia followed by fever were reported from on-line patient feedback, hence the display of temperature effects following vaccination was deemed to be fairly typical, although this case was particularly well characterized as the physical activity regimen was a precursor to a significant dynamic temperature response.

**Table 1:** The effective temperature change in response to vaccination and exercise over a three day period, of which the fourth day was a return to baseline. The response characteristics follow a 8:3:1 relative weighting among each day with the high temperature *ΔT*_fever_, low temperature *ΔT*_chill_, time periods *Δt*_chill_, *Δt*_fever_ and ramping periods completing the sequence 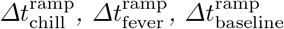.

| Day | $\Delta T_{\text{chill}}$ | $\Delta T_{\text{fever}}$ | $\Delta t_{\text{chill}}$ | $\Delta t_{\text{fever}}$ | $\Delta t_{\text{chill}}^{\text{ramp}}$ | $\Delta t_{\text{fever}}^{\text{ramp}}$ | $\Delta t_{\text{baseline}}^{\text{ramp}}$ |
| --- | --- | --- | --- | --- | --- | --- | --- |
| 1 | $-0.62^\circ\text{F}$ | $3.2^\circ\text{F}$ | 35 min | 150 min | 25 min | 60 min | 120 min |
| 2 | $-0.23^\circ\text{F}$ | $1.1^\circ\text{F}$ | 20 min | 60 min | 70 min | 120 min | 120 min |
| 3 | $-0.08^\circ\text{F}$ | $0.4^\circ\text{F}$ | 60 min | 30 min | 70 min | 120 min | 140 min |

The motivation for implementing this procedure to study temperature dynamics offers several advantages: (1) Model identification has been successfully deployed by the implementation of so-called step-responses, in which a step excitation in the input is made, and the output dynamics are studied: a model is then fit to the data, for example see the textbook on process control in the chemical industries [12]. The use of the data immediately after completing a session of physical activity can therefore be modeled as a step response in the input *E*(*s*) and the time constants can be clearly identified. (2) Because the exercise session was the normal routine, which had been repeated in previous days, at the same time, and under essentially the same *a priori* conditions, a clear response due to the impact of the vaccination could be realized. Since there were no changes to the normal routine, other than the vaccine injection, there was no other explanation for the trigger but the vaccine. (3) To one not engaged in daily physical activity, the protocol may seem rigorous; however, if it is routinely performed on a daily basis, it can be performed without much effort, and thus effects due to over-exertion could be ruled out as causative. (4) The response of “chill” followed by fever is hypothesized in this research to be due to a cardiovascular signal not in-synchronization as usual with the neural signal associated with the temperature. Hence measuring oral temperature would not be a good indicator to track the response. It has to be performed in which the patient can provide indication of the effects such as shivering effects in order to quantify the response, since it was induced by cardiovascular effects and not merely temperature effects. Thus, through patient interview, a quantitative assessment of temperature dynamics based on prior experience can be achieved. (5) By repeating the experiment over a multi-day period, the effect of lowered concentration of vaccine can be examined on the impact of the dynamic modes associated with the temperature response.

To quantify this response and evaluate the model and feedback control law, an effective temperature change trajectory was generated for each of the three days during the period of interest. The trajectories in which the effective temperature is quantified as based on prior experiences, are relative to each from day to day based on the intensity ratio of 8:3:1. It is noted that the response is one in which the effective temperature that is induced actually is modeled as dropping in magnitude to reflect the experiences of the patient, and then followed by a rise to a fever pitch consistent with prior experiences, followed by a breaking of the fever via sweating with a return to baseline. While the actual temperature need not be measured, thus the data is considered to be an “effective temperature” and the key points of the analysis are the relative departure in amplitude from nominal conditions on multiple sequential days and the precise timing associated with the temperature trajectory events. Since the participant performed physical activity for multiple days prior to the vaccination, the amplitude and time response from nominal could be well characterized for the routine. Thus, the model is scaled to reflect deviations from nominal conditions in which the time constants and associated dynamic characterization will be accurate and consistent with the model derivation. From this set of data points, a linear spline function is fit to the data points, which is then used as input to the system identification algorithm. The time scale for all analysis is performed in minutes, including the subsequent temporal and spectral characterization. The initial values of 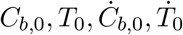 are selected as steadystate just prior to completing the physical activity routine, and thus the excitation input to the system is the step transition from physical activity to no physical activity, which enters the model through *E*(*s*).

To evaluate the ability of the model to capture the characteristic temperature response, a model for each of the three days is correlated to the data using system identification techniques [13] available in the Matlab programming language with the forcing function of a step-down in *E*_1_(*s*) considered as the exogenous input of physical activity. As consistent with the model derivation, the model order was selected as four poles and three zeros, and the mean square error was minimized in determining the model parameters. Using the multiple, linear time-invariant model representation about each new operating point, a model was produced for each of the three days. The temperature responses were then plotted in comparison to the data as presented in Figure 6, which show agreement in mapping the inverse response corresponding to hypothermic signs of shivering, followed by the temperature rise, either plateauing or momentarily hitting a peak as fever, and then terminating from a cool-down to baseline temperature through sweating.

**Figure 6:**
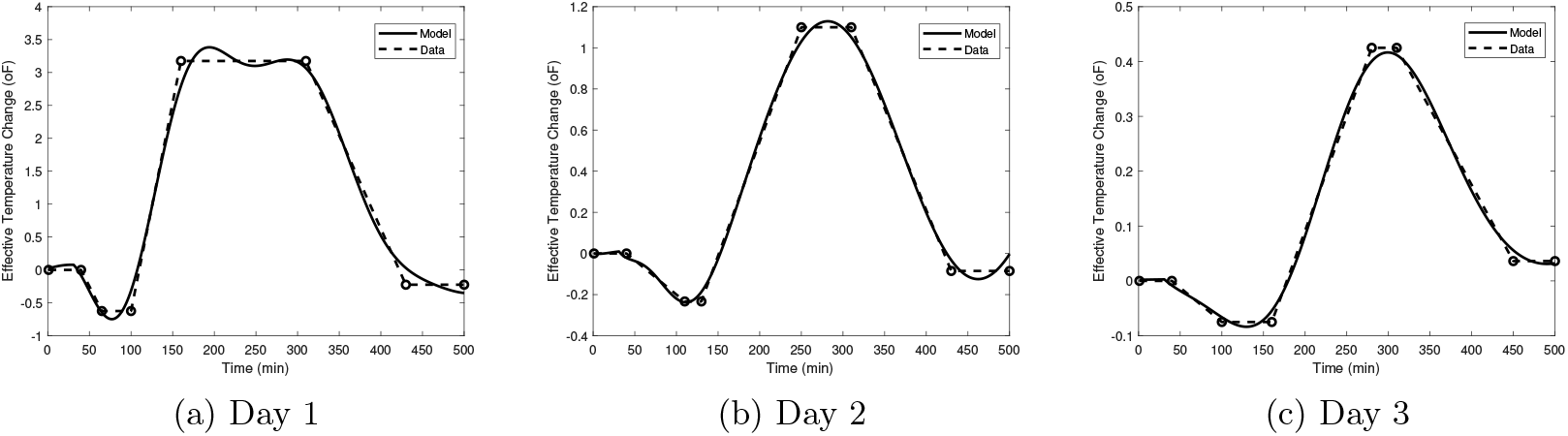
As an initial indication of model validation, a comparison of the measured and modeled trajectory of temperature is compared for each of the three days. The inverse dynamics of the model are clearly captured in showing a dip down in temperature followed by a rise and then a cool down back to base. For Day 1, some oscillation at the level off period in temperature is noted, while the static gains in settling the response for each of the days is indicated. Thus, the relatively close representation of the model output to that observed would indicate that the modeling approach is appropriate and that it can be examined for predictive and analysis capability.

The models in Laplace transform format of the closed-loop system response for each of the three days is listed in Table 2 including a goodness of fit metric, which is the normalized root mean square criterion, a measure of the capability of the model to reproduce the observed data relative to the mean of the data. The goodness of fit exceeds 90% for each of the three models, which typically is required to produce acceptable correlation of the dynamic model to the data.

**Table 2:** The closed-loop models, whose structure order was derived by modeling, that map body temperature response to exercise, or activity, input, independent of the particular trajectory of input excitation. Thus, the temperature response to various scenarios can be predicted, and the internal dynamics of the body temperature control can be studied in response to differing concentrations of vaccines as it is cleared from the body. The goodness of fit parameter measures the ability to replicate the observed data relative to the mean of the data, with in exceeding 90% for each of the days indicating a good fit for system identification purposes.

| Day | Model Fit | Closed-Loop Model $G^c(s)$ |
| --- | --- | --- |
| 1 | 92.8% | $\frac{0.01276s^3+0.001238s^2-1.839e-05s-3.836e-07}{s^4+0.01148s^3+0.001893s^2+4.122e-06s+2.825e-07}$ |
| 2 | 95.4% | $\frac{0.005706s^3-0.0002954s^2+3.322e-05s-4.696e-07}{s^4+0.04177s^3+0.00365s^2+1.595e-05s+9.973e-07}$ |
| 3 | 94.8% | $\frac{0.001407s^3-2.516e-05s^2+3.173e-06s-3.735e-08}{s^4+0.01384s^3+0.001179s^2+6.037e-06s+2.242e-07}$ |

In assessment of the internal dynamics of the closed-loop models and the transition of daily trajectories while the effects of the vaccination are attenuated, the frequency response of the models for each of the three days is presented in Figure 7. For feedback control analysis, the frequency response is a useful indicator of the limits of performance [14]. It is noted that the second harmonic is strong on the first day, and then is less over the next two days. As the response curves tend to show a response closer to a single oscillatory function, one would expect a stronger magnitude at a single harmonic for Day 2 and Day 3, while on Day 1, the transition into a fever, which is held for some time, requires additional faster harmonic functions to model appropriately. The characteristics of the frequency response shows the static gain at low frequencies, and the resonant frequency indicates the peak response based upon the excitation of the input signal, which in this case is through physical activity.

**Figure 7:**
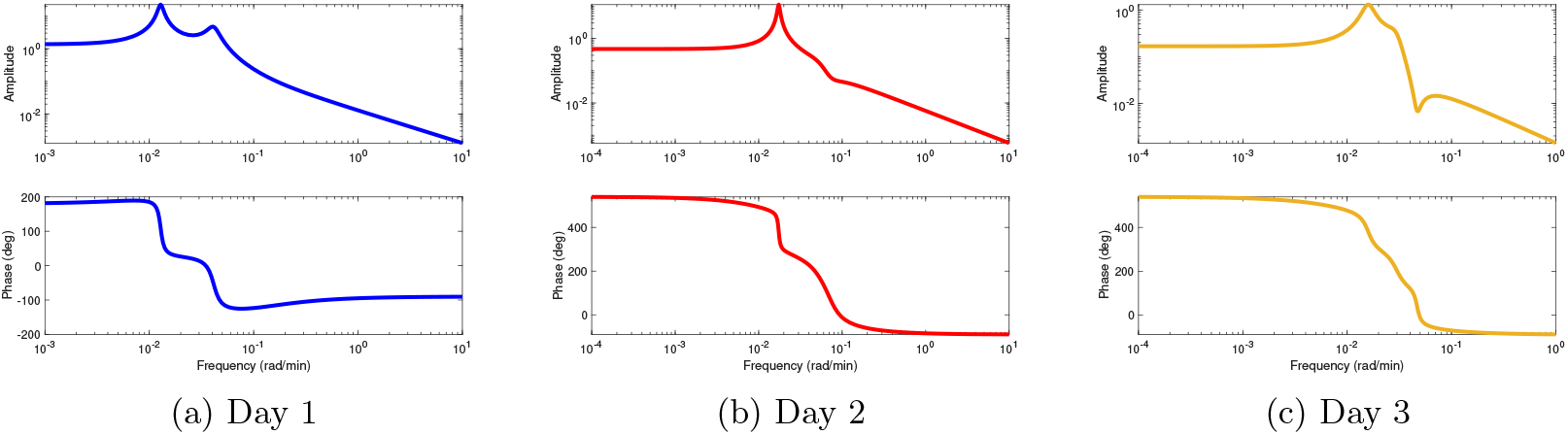
To characterize the types of responses in body temperature, the frequency response for each of the days is presented, which suggests that there are resonance modes that if matched or excited by the input, will result in significant deviation in temperature relative to other modes. This is useful as will be seen later in the paper on optimizing the input to the system in order to minimize fluctuation in temperature when infected. The shape of the curves are approximately the same from Day 1 to Days 2 and 3 as the vaccine is cleared, but the second resonant mode, which is associated with faster dynamic response, is reduced in amplitude, indicating a smoother trajectory represented by a single oscillatory state.

In Table 3, the location of the poles and zeros, as well as the gain margin and phase margin, of the identified closed-loop models is presented. It is noted that there is a right half plane zero for each of the models, which is indicative of the inverse response of chills and shivering preceding fever and subsequent sweating. The poles of the model are all negative, which is an indicative of stability, and the poles are of an imaginary number, indicating the oscillatory nature of the system response.

**Table 3:** The dynamic characteristics of the Closed-Loop models for Day 1, 2 and 3 is compared. For each of the models, there is a zero located in the right half plane, which indicates that it is non-minimum phase and for a step input the output trajectory will follow an inverse response, in which the effective temperature will drop before it rises, that is chills before fever. The poles are imaginary numbers, which indicates that the response will contain oscillatory response modes as it is closer to instability. The margins indicate are adequate for Days 2 and 3 indicating robustness to disturbances, but for Day 1, it indicates lack of margin.

| Day | Zeros | Poles | Gain Margin | Phase Margin |
| --- | --- | --- | --- | --- |
| 1 | -0.1078<br>0.0229<br>-0.0122 | $-0.0050 \pm 0.0411i$<br>$-0.0007 \pm 0.0128i$ | -2.66, -17.1 | 58.6 |
| 2 | $0.0181 \pm 0.0702i$<br>0.0157 | $-0.0203 \pm 0.0538i$<br>$-0.0006 \pm 0.0174i$ | 6.54, 14.4 | -52.4, 99.1 |
| 3 | $0.0029 \pm 0.0467i$<br>0.0121 | $-0.0049 \pm 0.0294i$<br>$-0.0020 \pm 0.0158i$ | 15.6, 13.1 | -113, 153 |

The Nyquist plot of the frequency response is presented in Figure 8 in comparing the dynamics of Day 1, Day 2, and Day 3, as the body transitions higher concentrations of the vaccine to lower concentrations as it is depleted. The vaccination is seen to have a significant effect on the dynamic response. Clearly, while the vaccination agents are active the temperature dynamics are compromised as reflected in the ratio of intensity of the temperature response comprising the data.

**Figure 8:**
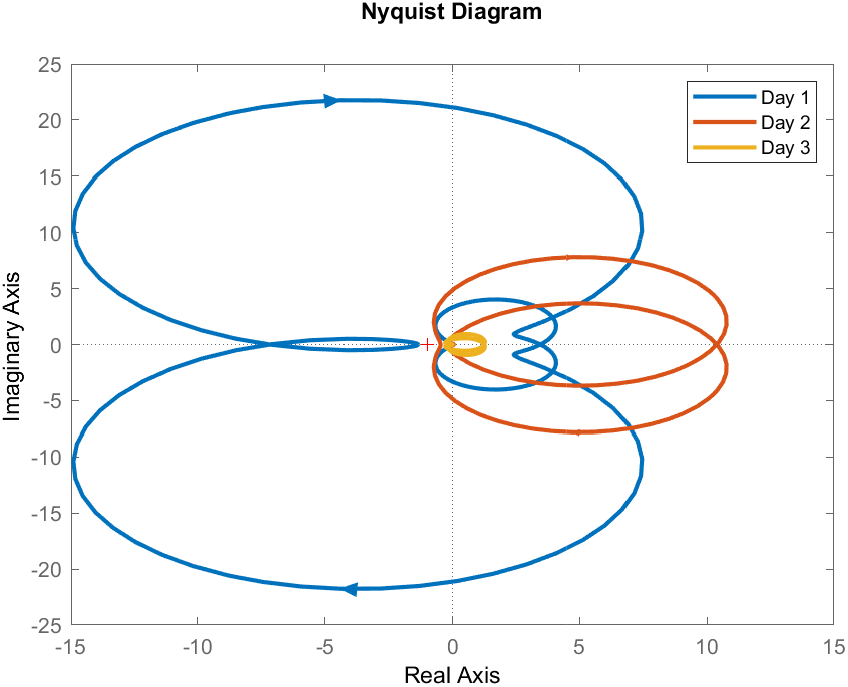
In evaluating the impact of the vaccine on body temperature control, the Nyquist representation of the models for Day 1, 2, and 3 is compared to show dramatically the severe degradation of temperature control due to the influence of the polysaccharide on the internal dynamics. The Nyquist plot shown an encirclement of the (−1, 0), which indicates instability if a secondary feedback controller were to be wrapped around the intrinsic closed-loop temperature control system. There is a significant reduction in the representation in Day 2 and in returning close to nominal behavior of Day 3.

#### 3.8.1 Verifying Control Law Structure

To assess the assumptions used in deriving the control law and subsequent modeling equations, it is useful to evaluate the model error predicted by the derivation and its ability to fit the data accurately. If the model is overparameterized, then lower order models would be able to fit the data, while if the model is under-parameterized, for example if critical modes are neglected, then higher order models will fit the data dramatically better. As indicated, the model derivation indicates that the temperature dynamics in response to the step function can be represented as 4 poles and 3 zeros at a nominal operating point. Investigating the model order therefore as a means of evaluating the control law hypothesis, the mean square error of different model orders is presented in Table 4 for each of the trajectories. The (4,3) model performs roughly 7 times better than the (3,2) model, while the (5,4) model is only marginally better at 20% fit; thereby indicating that the (4,3) model is an adequate representation of the data. This trend is the same for Day 2 and Day 3.

**Table 4:** For a validation of the model structure, the mean square error of the fitted parameter models are compared to the 4 pole, 3 zero structure that was derived. An over parameterized model of 5 poles, 4 zeros, and an under parameterized model of 3 poles, 2 zeros is shown, which indicates that the derived structure provides about 7X enhancement on Day 1 over the under parameterized model, while providing essentially the same predictive quality as the over-parameterized model, thereby indicating an appropriate structure, and some evidence of validation of the modeling approach. Days 2 and 3 produce similar results, although less dramatic since the vaccine is being cleared from the body and thereby inducing less oscillation and transient deviation from nominal.

| Day | (3,2) Model | (4,3) Model | (5,4) Model |
| --- | --- | --- | --- |
| 1 | $8.9 \times 10^{-2}$ | $1.1 \times 10^{-2}$ | $9.1 \times 10^{-3}$ |
| 2 | $7.1 \times 10^{-4}$ | $4.7 \times 10^{-4}$ | $2.7 \times 10^{-4}$ |
| 3 | $1.6 \times 10^{-4}$ | $7.4 \times 10^{-5}$ | $1.1 \times 10^{-4}$ |

As an example of the model fitting capability, the confidence band of the standard deviation surrounding the model response for Day 2 is depicted in Figure 9. It indicates that the model does well in capturing the key points of the response which are the breaking points and times associated with the dynamic trajectories of the hypothermic phase followed by a heating to a fever pitch and a return to baseline by sweating.

**Figure 9:**
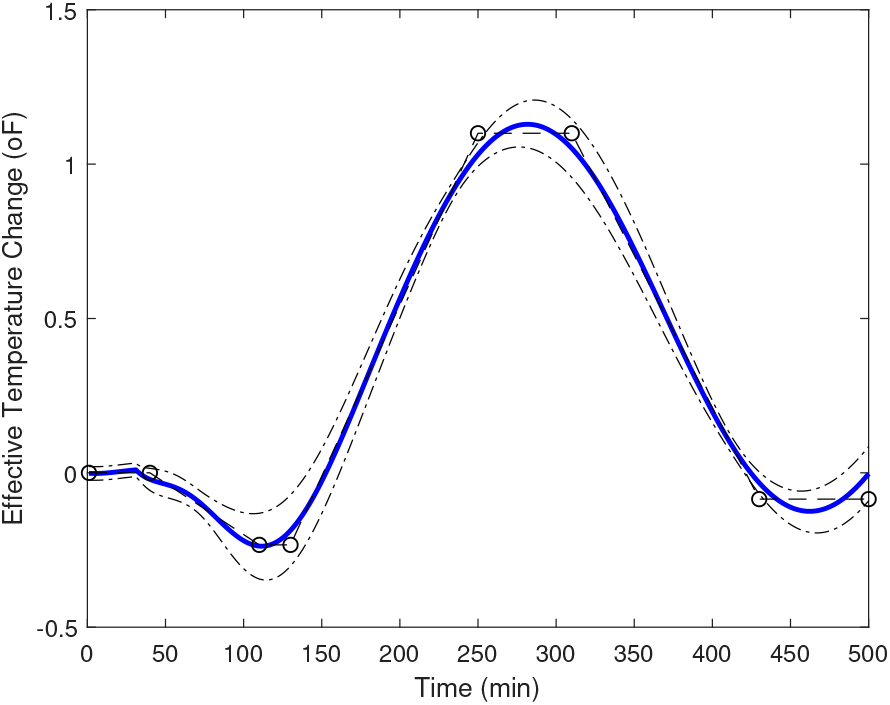
The standard deviation bound is shown for the temperature response of Day 2, indicating tight fitting along the trajectory up and down, with looser fitting during the initial phase of the response and the settling value.

To further examine the adequate representation of the system dynamics by the (4,3) model predicted by the derivation, the consequence of the model order is compared in Figure 10 for the fit of Data to the Day 1 trajectory for a model with 3 poles and 2 zeros, and another model with 5 poles and 4 zeros. As seen from the result, the (3,2) model fails to capture an important node of the response, showing a single oscillation, and the (5,4) model falls almost exactly on the (4,3) model, suggesting that the addition of an extra pole and extra zero does not improve the predictive capability. Consequently, it is concluded that a (4,3) model is appropriate model to relate the observed temperature response when subjected to exogenous input in the presence of infections that alter the fundamental dynamic modes.

**Figure 10:**
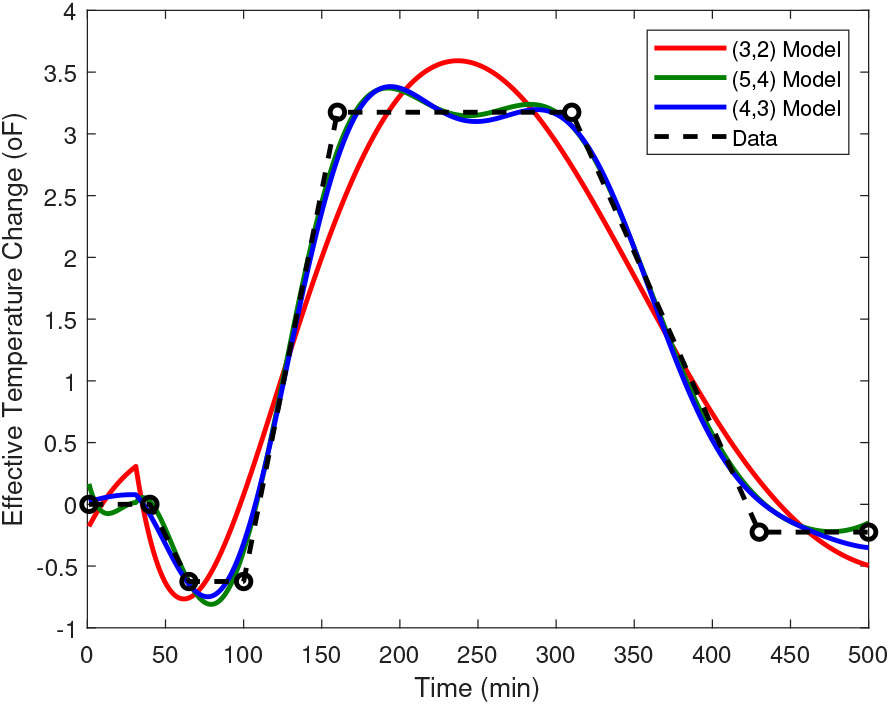
To validate the model structure, a comparison of model order is conducted to see that the data is neither over or under parameterized, which would indicate that response modes are either extraneous or neglected. The comparison is to the 4 poles, 3 zeros model predicted by the derivation to models of the 3 poles, 2 zeros and 5 poles, 4 zeros. The results indicate that 4 poles, 3 zeros is appropriate as characterized by the dynamic trajectory of the first day, which basically coincides with the higher order model, while providing significantly more information than the lower order model, which apparently neglects one oscillatory mode.

#### 3.8.2 Influence of infectious agent on temperature dynamics

As noted from the analysis of the transient and spectral characteristics of the identified models for Days 1, 2, and 3, the vaccination has a significant influence on the temperature response when stressed, and the fundamental nature proposed is the interference of PGE2 with its interaction with cortisol at the LBD of GR. Further, as noted in Figure 11, which represents a relation of the differences of the amplitude and phase between the response associated with Days 1 and 2, the differences are due not only to the reaction kinetics of cortisol but also the dynamics associated with the PGE2 depletion via dissipation of the vaccination molecules. Had it only been due to the cortisol reaction rate, the difference in model structure would have been a constant amplitude and a phase associated with the time delay; however, the response is more complex indicating the participation of each element associated with the interaction between PGE2 and cortisol. The transition from Day 1 to Day 2 for the two LTI representations, has break points at 0.01 rad/min and 0.04 rad/min, corresponding to time constants of 100 minutes and 25 minutes, or roughly 5 hour and 1 hour response periods.

**Figure 11:**
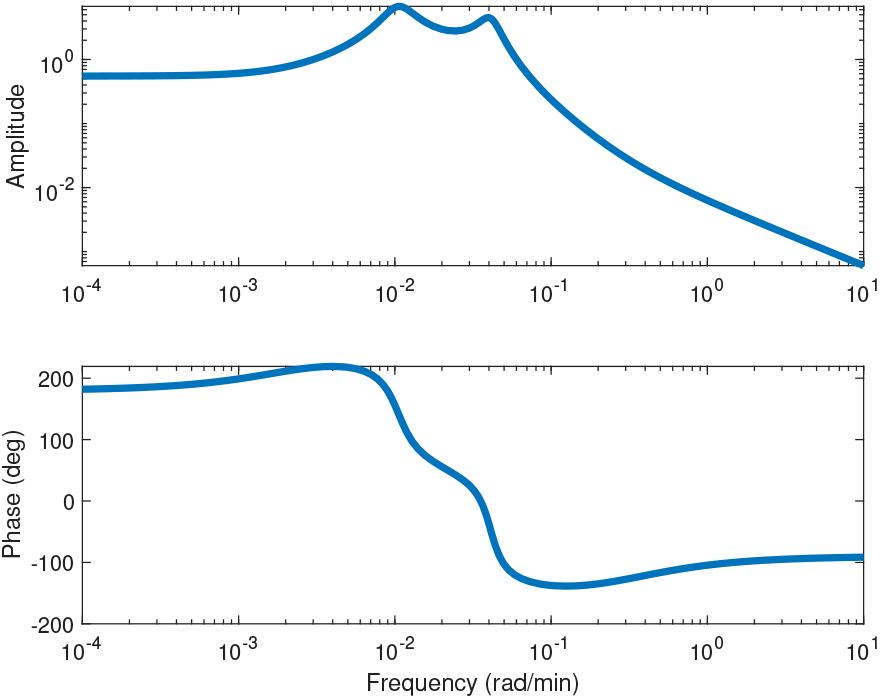
To assess the transition in model dynamics as the vaccine is cleared from the body, the difference of the frequency response is examined for the models of Day 1 and Day 2, which indicates that the resonant peaks at time constants of approximately 100 minutes and 25 minutes, resulting in expected in how the body temperature will respond in both short term and long term, roughly 5 hour and 1 hour response periods.

#### 3.8.3 Minimizing Temperature Fluctuations

One question that can be assessed using the model response is whether the observed temperature fluctuations of the case study due to the abrupt cessation of exercise would have been reduced had there been an extended cool down period in which the exercise was gradually reduced. This would seem to be the case since the frequency response of the closed-loop model indicates certain peaks which if excited would result in larger oscillatory responses, and therefore if the excitation can be kept to low frequencies, then less peak temperature fluctuation should be expected. This turns out to be the case, as seen in Figure 12, which compares the temperature fluctuation observed for Day 2, when the exercise was immediately stopped, in comparison to the temperature fluctuation had the exercise been gradually ramped down for the full period of observed nominal temperature oscillation. It is noted that temperature deviations during the period of physical activity were not observed because it was slow relative to the cessation of the physical activity, and hence essentially a ramp, which can be modeled as 1/*s*^2^ that has significantly more frequency roll-off and attenuation then the step response of 1/*s*.

**Figure 12:**
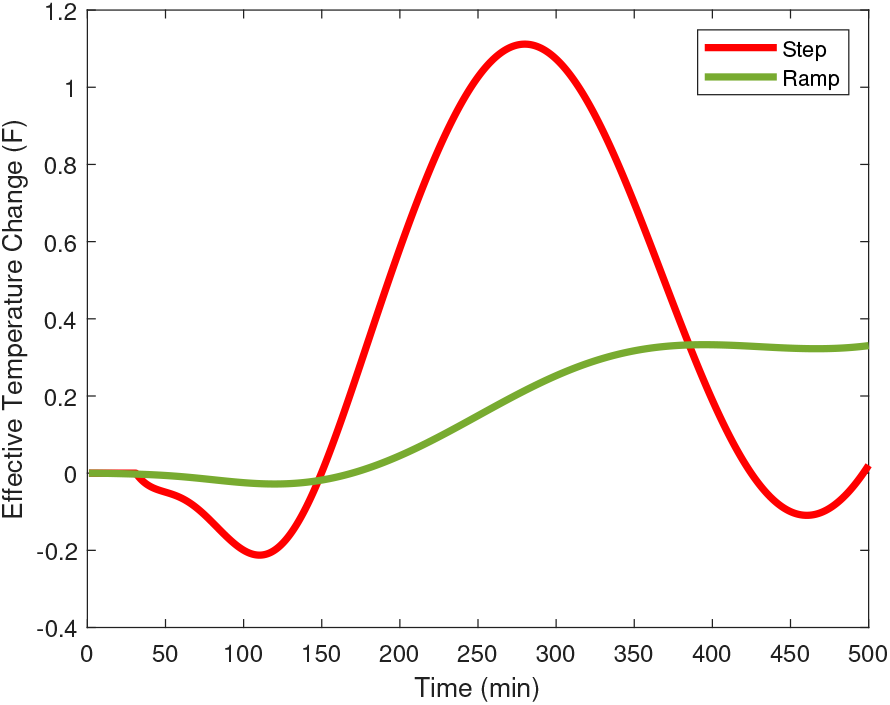
The model has predictive capability in optimizing and planning voluntary activity to compensate for sub-optimal situations. In this example, the temperature deviations are minimized if the cool down period after exercise were to have been conducted over an extended period, represented as a slow ramp down in activity, relative to just stopping the exercise regime, represented as step-down in the case study. The slow ramp down avoids the excitation of resonant frequencies in the dynamic response, which are present due to the vaccination altering the inherent dynamic response, pushing the system closer to instability in inducing oscillatory temperature modes.

## 4 Discussion

As verified through the multivariable model and experimental response study, the reasoning for the inducement of fever travels through the association of PGE2 with the ligand binding domain of glucocorticoid receptors. Nominally, cortisol occupies the LBD space, and the insertion of PGE2 at the LBD prevents the expression of cortisol according to its designed and expected performance. The accumulation of cortisol within the GR and the low conversion rates alter the passage rate of the hormone through the plasma membrane, and the additional incorporation of the calcium ion, changing its permissive nature, and altering the firing rates of the neurotransmitter induced thermal effectors. Thus, PGE2, which experimentally has been noted to be a pyrogen, is ultimately responsible for the temperature response of chills and fever.

In addition to PGE2, cortisol is also central molecule to the dysfunctional temperature responses. It interacts at the cellular level in both the decision making center, which implements the commands, and also at the target tissues to enhance the sensitivity of the cell through its localized depolarization induced by passage through the plasma membrane, and carrying two calcium ions. Cortisol is characteristic of the status of the cardiovascular system, and that information is taken into account through the temperature control commands. Moreover, in addition to its concentration, the rate at which cortisol also changes is important to autonomic control of temperature.

The dynamics of temperature are characterized as a linear set of equations, whose modes or poles, are a function of the concentration and rate of cortisol, and thus are subject to the concentration and rate of incorporation of PGE2. The shift in the dynamics due to the presence of the agents may result in strange responses, such as an inverse response, that is chills before fever, and oscillatory behavior, which is primarily responsible for fever. The time delays in the system, mainly from transportation delays, but also from the inherent slow dynamics of cooling, will limit the performance of the control system. Because the internal dynamics of temperature control are compromised during infection, temperature effects can be minimized by avoiding significant excitations to it, such as load disturbances.

The results are consistent with the action observed of fever in dysfunctional or diseased states. For example, PGE2 is a known pyrogen, and this research article indicates its reason behind it. Moreover, if cortisol levels run very low, adrenal insufficiency results, with signs including fever[15], which is even at times misdiagnosed as an infection [16]. Further correlations can be inferred from the undulation of fever characterizing disease such as brucella [17] and the rise and fall of CORT during the day and night [18] as does body temperature.

In addition, it is noted that injections of hydrocortisone can control a fever [19], which is consistent with this study. With the addition of glucocorticoids, the fever is actually controlled by the diluting effects on PGE2 in a displacement at the GR, thereby allowing CORT or a glucocorticoid to be processed effectively. As noted PGE2 is a more stable configuration with the LBD, and thus a higher concentration of glucocorticoids will be necessary to shift the equilibrium away from PGE2, allowing time for its ultimate removal or degradation. Further, it is indicated that the mathematical models can be used to optimize the release of corticosteroids in order to avoid unnecessarily high levels within the cell.

In addition to the identification of its underlying molecular cause, this research article also provides insight into the nature of a fever. From the analysis, it is clear that it is not necessary to define a “setpoint” but rather an operating mean from which temperature is regulated. As a defense mechanism, fever may not control significantly the rates of association with the LBD of PGE2, since a differential would have to be established with cortisol. Because of their close molecular weight it would not seem substantial, however the linear framework at a higher temperature may be preferred, which would lower the rate of conformation in order to achieve a low energy state with the LBD. These results extend to the nucleus and the basic characterization and interaction of DNA and steroid hormones, as described in [20].

It was interesting that a vaccine could produce a fever in such a straightforward manner, merely involving a sustained period of physical activity, followed by an abrupt cessation. However, it is noted that the cortisol levels run high during prolonger physical activity and thus it is to be expected, and should be indicated during the injection of the vaccine. The pathway involved the increase concentration of cyclooxygenase, and thus was not as significant, as if it were to generate arachidonic acid, but still the conversion into PGE2 made it indistinguishable from a nominal fever.

Another interesting facet of this work was the methodology in which a mathematical model of the response was developed before the chemistry was identified. Thus, this adds to the credibility of the approach and to the model, because it is consistent with the underlying performance, and not merely a curve fitting routine. Perhaps other diseases or unknowns in biology can be resolved through a similar approach, since the mathematics can be used to significantly confine the molecules that could satisfy the basic operational equations. In addition, it highlights the complexities in that it is necessary to consider the mathematical characteristics of the interactions to identify the root causes, as mere correlation studies will not explain the phenomena: for example, the rate and interactions are important.

Following this approach, other signs and symptoms can be modeled and understood from a first principles basis, as it is my belief that the inhibition of prostaglandins with cortisol at the LBD are responsible for a great many dysfunctional responses, including energy related responses, such as fatigue and malaise, as well as cerebral effects, such as fogginess and depression, during diseased states. It may also be noted that the interference at the GR due to the similarity of prostaglandins and corticosteroids may also induce diseased effects involved cardiac and renal diseases, including impact on the similar aldosterone. It seems clear that the tightly coupled dynamics of multiple trajectories and diseases could be induced by the dysfunction of just a single dysfunctional associations, brought on and accelerated by the presence of infectious agents or those factors associated with the inflammatory response.

## 5 Methods

- Molecular Modeling: The software system called Avogadro was used. The ligand binding domain residues, which were indicated in [21], were approximately positioned relative to CORT, and an optimization was run with the MMFF94 [22] force field to position the elements and calculate the bond lengths so as to minimize energy state. CORT was then removed and replaced with the conformer of PGE2, and the optimization routine was rerun to optimize the position of the elements and bond lengths in order to minimize the energy level. Calcium ions were added at the areas of high negative electrostatic potential, as described in [5].
- Experimental: A participant’s regularly scheduled vaccination via pneumococcal agent (Pneumovax 23) was administered by a physician followed by a blood test at an accredited medical services institution. An influenza vaccination (Flublok Quadrivalent) was given at the same time. Six hours after vaccination, physical activity was performed consisting of a two hour work effort, including ninety minutes of treadmill at moderate speed average 4.7 miles per hour. Perceived temperature was monitored over time and relative temperatures were recorded afterwards at approximately fifteen minutes for the first three hours, followed by approximate hourly assessment. This same procedure in physical activity was performed for the following three days, with the same physical activity protocol utilized. Fatigue was also monitored, noting that the physical activity achievable was less than the prior day of vaccination. This procedure was useful in that it could trigger a significant likelihood of body temperature fluctuations as it induced high levels of cortisol through physical stress over an extended period, high levels of PGE2 through the injection of the vaccination with a wait period allowing for distribution as it involved the TNFα pathway [23], and a high load placed on the internal control system by immediately stopping physical activity thereby exciting the body temperature control system.
- Identification of thermoregulation model: To evaluate the ability of the model to produce the characteristic temperature response, a model for each of the three days was determined using system identification techniques using the Matlab programming language, system identification toolbox [13]. A forcing function of a step-down in physical activity was considered as the exogenous input of physical activity. As consistent with the model derivation, the model order was selected as four poles and three zeros, and the mean square error was minimized in determining the model parameters. The modeling parameters were used to minimize the mean square error of simulation.

## Bibliography

[1] J. E. Hall, Guyton and Hall textbook of medical physiology. Elsevier Health Sciences, 2015.

[2] Ask the Experts, “What causes fever,” Scientific American, no. 2, 2006.

[3] A. A. Romanovsky, “The thermoregulation system and how it works,” in Handbook of Clinical Neurology. Elsevier, 2018, vol. 156, pp. 3–43.

[4] C. Schaper, “Dynamics and Internal Control of Body Temperature in Response to Infectious Agents and Other Causal Factors,” bioRxiv, 2019, https://doi.org/10.1101/566679v1.

[5] “Competitive Inhibition of Cortisol by Prostaglandins at the Ligand Binding Domain of Glucocorticoid Receptors,” bioRxiv, 2019, https://doi.org/10.1101/851501.

[6] N. S. Kirkby, M. V. Chan, A. K. Zaiss, E. Garcia-Vaz, J. Jiao, L. M. Berglund, E. F. Verdu, B. Ahmetaj-Shala, J. L. Wallace, H. R. Herschman et al., “Systematic study of constitutive cyclooxygenase-2 expression: role of nf-κb and nfat transcriptional pathways,” Proceedings of the National Academy of Sciences, vol. 113, no. 2, pp. 434–439, 2016.

[7] J.-l. Lai, Y.-h. Liu, C. Liu, M.-p. Qi, R.-n. Liu, X.-f. Zhu, Q.-g. Zhou, Y.-y. Chen, A.-z. Guo, and C.-m. Hu, “Indirubin inhibits lps-induced inflammation via tlr4 abrogation mediated by the nf-kb and mapk signaling pathways,” Inflammation, vol. 40, no. 1, pp. 1–12, 2017.

[8] J. Chetsawang, S. Nudmamud-Thanoi, R. Phonchai, Z. Abubakar, P. Govitrapong, and B. Chetsawang, “Methamphetamine toxicity-induced calcineurin activation, nuclear translocation of nuclear factor of activated t-cells and elevation of cyclooxygenase 2 levels are averted by calpastatin overexpression in neuroblastoma sh-sy5y cells,” Neurotoxicology, vol. 67, pp. 287–295, 2018.

[9] S.-H. Kim, J. Roszik, S.-N. Cho, D. Ogata, D. R. Milton, W. Peng, D. G. Menter, S. Ekmekcioglu, and E. A. Grimm, “The cox2 effector microsomal pge2 synthase 1 is a regulator of immunosuppression in cutaneous melanoma,” Clinical Cancer Research, vol. 25, no. 5, pp. 1650–1663, 2019.

[10] G. Stephanopoulos, Chemical process control. Prentice-Hall Englewood Cliffs, NJ, 1984.

[11] A. Behrouzvaziri, “Thermoregulatory effects of psychostimulants and exercise: Data-driven modeling and analysis,” Ph.D. dissertation, 2018.

[12] D. E. Seborg, D. A. Mellichamp, T. F. Edgar, and F. J. Doyle III, Process dynamics and control. John Wiley & Sons, 2010.

[13] L. Ljung, System identification: theory for the user. Prentice-hall, 1987.

[14] S. Boyd and C. Barratt, “Linear controller design: limits of performance,” Stanford University Stanford, CA, Tech. Rep., 1991.

[15] R. L. Rushworth, D. J. Torpy, and H. Falhammar, “Adrenal crisis,” New England Journal of Medicine, vol. 381, no. 9, pp. 852–861, 2019.

[16] R. Dineen, C. J. Thompson, and M. Sherlock, “Adrenal crisis: prevention and management in adult patients,” Therapeutic advances in endocrinology and metabolism, vol. 10, 2019.

[17] C. G. Ephrem, M. J. Matar, G. C. Chalhoub, and W. G. Greige, “Brucella endocarditis: diagnostic challenges.” Journal of Infection in Developing Countries, vol. 12, 2018.

[18] S. S. Simons, R. Beijers, A. H. Cillessen, and C. de Weerth, “Development of the cortisol circadian rhythm in the light of stress early in life,” Psychoneuroendocrinology, vol. 62, pp. 292–300, 2015.

[19] A. Cilliers, A. J. Adler, and H. Saloojee, “Anti-inflammatory treatment for carditis in acute rheumatic fever,” Cochrane Database of Systematic Reviews, no. 5, 2015.

[20] C. Schaper, Design of DNA, Genetic Codes, and Life Function. ISBN 978-1-73537210-5. Molecular Sciences Publishing House, 2020.

[21] X. Liu, Y. Wang, and E. A. Ortlund, “First high-resolution crystal structures of the glucocorticoid receptor ligand-binding domain–peroxisome proliferator-activated γ coactivator 1-α complex with endogenous and synthetic glucocorticoids,” Molecular Pharmacology, vol. 96, no. 4, pp. 408–417, 2019.

[22] T. A. Halgren, “Merck molecular force field. i. basis, form, scope, parameterization, and performance of MMFF94,” Journal of computational chemistry, vol. 17, no. 5-6, pp. 490–519, 1996.

[23] O. Elkayam, D. Caspi, T. Reitblatt, D. Charboneau, and J. B. Rubins, “The effect of tumor necrosis factor blockade on the response to pneumococcal vaccination in patients with rheumatoid arthritis and ankylosing spondylitis,” in Seminars in arthritis and rheumatism, vol. 33, no. 4. Elsevier, 2004, pp. 283–288.

